# Oblique plane single-molecule localization microscopy for thick samples

**DOI:** 10.1101/289686

**Authors:** Jeongmin Kim, Michal Wojcik, Yuan Wang, Ke Xu, Xiang Zhang

## Abstract

We introduce single-molecule oblique plane microscopy (obSTORM) to directly image oblique sections of thick samples into depth without lengthy axial stack acquisition. Using oblique light-sheet illumination and oblique fluorescence detection, obSTORM offers uniform super-resolution throughout imaging depth in diverse biological specimens from cells to tissues. In particular, we demonstrate an isotropic resolution of ∼51 nm over a depth of 32 μm for a tissue sample, and comparable resolution over a depth of 100 μm using fluorescent beads.

---

Single-molecule localization microscopy (SMLM)^1-3^ has become an important tool for biology thanks to its superior resolving power. Typical SMLM setups capture lateral plane (*xy*) images with added axial (*z*) information^4-6^ over a shallow range of ∼1 µm. Extended views through the sample depth may be extracted indirectly by stitching axial stacks of three-dimensional (3D) SMLM images^7^, but this approach is extremely slow, error-prone, and susceptible to photobleaching. Moreover, the accessible imaging depth of SMLM is often limited to a few microns from the coverslip surface due to the intrinsically shallow depths of illumination and detection associated with the use of oil-immersion objectives for confined illumination by total internal reflection or near-critical angle of incidence^8^. Although selective plane (or light-sheet) illumination has enabled deeper SMLM imaging^9-12^, samples often have to be prepared in specialized geometry and devices, and the achieved resolution is still suboptimal. Furthermore, to obtain depth information in oblique and axial views still necessitates difficult z-stack acquisition.

Here we report oblique plane super-resolution microscopy (obSTORM), a sectioning-free SMLM technique that directly images any oblique (including axial) plane through thick biological samples on regular glass coverslips. An oblique or axial view is achieved by optically conjugating the target oblique plane in a sample with a camera sensor plane^13-17^, so that the wide-field section at an oblique angle of α is imaged by a single-shot (**Fig. 1a**). While earlier, diffraction-limited oblique plane microscopy (OPM) was at small oblique angles (α ≤ 32°)^13-15^, we recently extended α to 90° in oil-immersion systems^17^. To enable single-molecule imaging through thick biological samples, we have redesigned the OPM system with a water immersion objective (O1) and engineered a tunable α of 45-90° (**Supplementary Fig. 1**). Moreover, we substantially improved light efficiency and reduced undesired ellipticity of point spread function (PSF) through the integration of a polarized beam splitter (PBS) and an achromatic quarter-wave plate (QWP) together with a remote objective (O2) of a larger cone angle than that of O1. (**Supplementary Fig. 2 and 3, Supplementary Derivation**). Samples for obSTORM are simply prepared on coverslips, and thus samples from cellular to tissue levels are easily accommodated.

**Figure 1.**
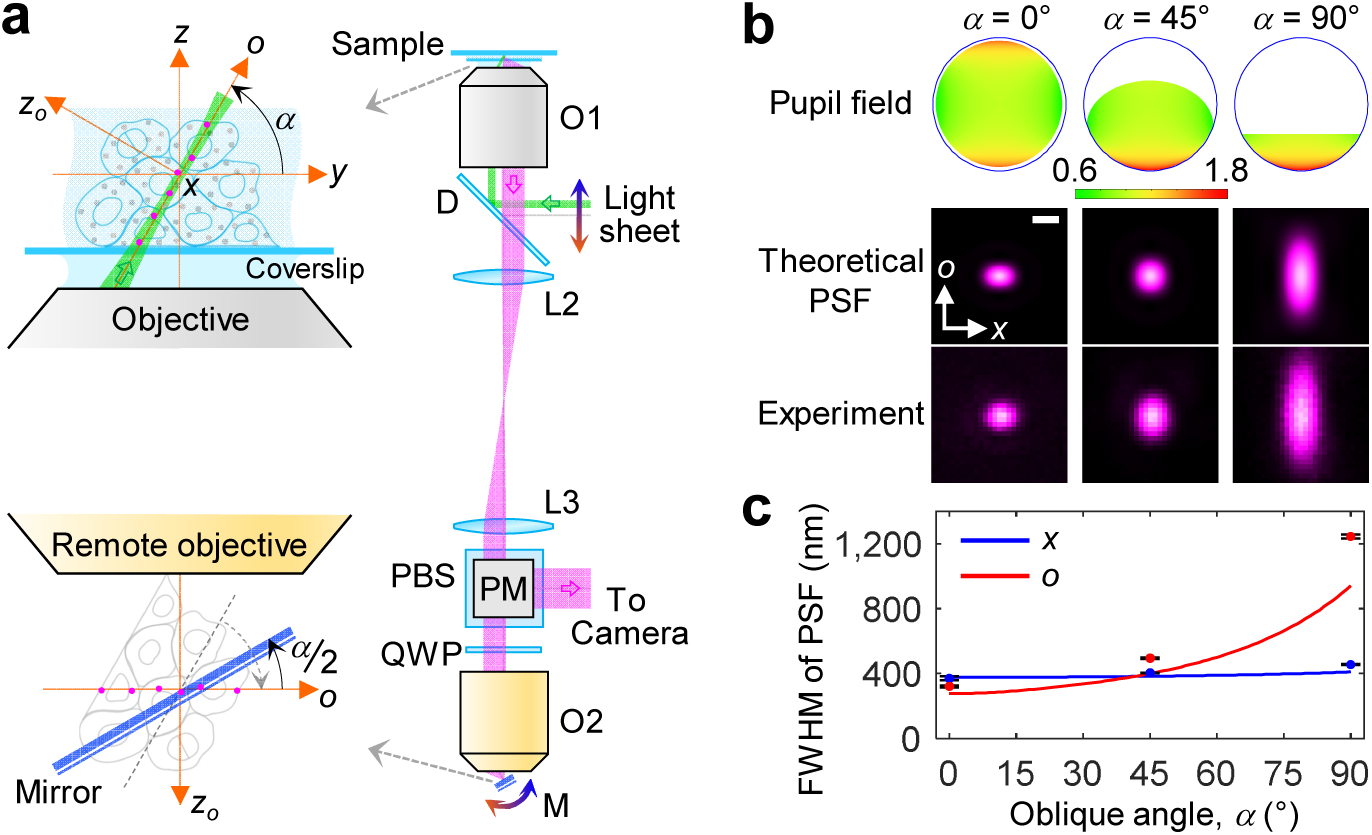
Imaging principle and PSF of obSTORM. (**a**) Schematic layout. An inclined light-sheet along the *xo* plane, rotated by (tunable) α from the focal plane of a water-immersion objective (O1, NA = 1.2), excites and activates single molecules through depth in a fluorescent sample. The sample space is isotropically mapped into the remote space of the second objective (O2, NA = 0.95) and then reoriented as illustrated by cell aggregate due to an oblique mirror (M) tuned at α/2. Thus, the single molecules emitted on the oblique plane (magenta dots) are conjugated to the focal plane of O2 and thus directly detected by a camera. A polarizing beam splitter (PBS) is oriented to transmit only *x*-polarized fluorescence signals from the sample. The PBS in combination with an achromatic quarter-wave plate (QWP) reflects the backward signals from O2 to the camera with no further loss. O, objective; D, dichroic beam splitter; L, lens; PM, periscope mirror. (**b**) Rotationally asymmetric exit pupil and electric field distribution (scaled to one at the pupil center), and resultant PSFs for 1.2/0.95 NA combination. A light clip at O2 by the oblique mirror decreases an effective pupil area by 30% and 70% at α = 45° and 90°, respectively. Stronger edge fields in the *o* direction shaped by the PBS mitigate the ellipticity of PSFs at higher oblique angles. PSFs were calculated at 685 nm wavelength and measured with dark-red fluorescent beads. Scale bar, 400 nm. (**c**) Comparison of theoretical (lines) FWHM of PSF over oblique angle with experimental results (dots and error bars; *n* = 9, 4 and 6 beads for 0°, 45°, and 90° respectively; error bars, s.d.).

We first characterized the PSF of the obSTORM system using fluorescent beads. Our theoretical calculations and experimental measurements both indicated near-circular PSFs for α = 0-50° with a modest increase in full-width at half-maximum (FWHM) to ∼400 nm when compared to that of typical conventional in-plane microscopy (∼280 nm), and this value is further increased in the vertical direction for higher angles (**Fig. 1b,c** and **Supplementary Fig. 4** for emission at 685 and 570 nm, respectively). Nonetheless, the PSF maintained its brightness and size over an axial depth of >100 μm with a uniform (within ∼15%) localization precision at α = 45° (**Supplementary Fig. 5**).

For Alexa Fluor 647 (AF647)-labeled microtubules in fixed A549 cells, obSTORM directly resolved the microtubule network throughout the whole cell depth along the 45° diagonal direction (**Fig. 2a,b**). The light efficiency of single molecules in such 45° obSTORM is approximately 25% of the conventional STORM efficiency owing to the following signal losses: 50% at the PBS, 26% at the remote objective (double pass), and 30% at asymmetric pupil (**Fig. 1b**). Thus, the average number of collected photons for each AF647 molecule per frame was 954 (**Fig. 2c**). This value is comparable to what is typically found in SMLM experiments based on fluorescent proteins (e.g., mEos) and none-AF647 dyes (e.g., Atto 488 and Nile Red), and single molecules images were obtained with high contrast (**Fig. 2b** inset and **Supplementary Video 1** at 50 frames per second).

**Figure 2.**
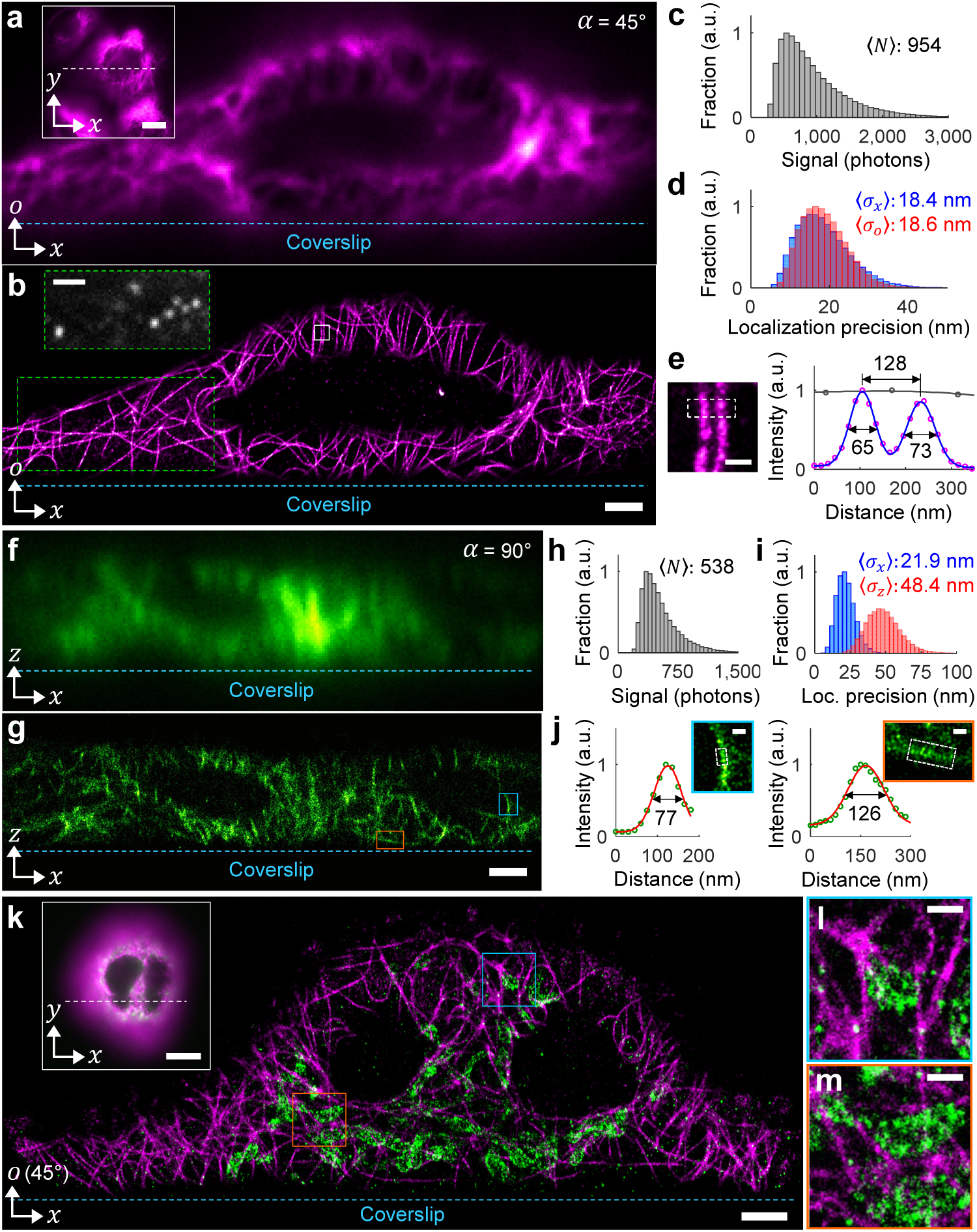
Super-resolution oblique plane imaging by obSTORM with A549 cells cultured on coverslips. (**a**) Diffraction-limited, 45° oblique plane fluorescence image of microtubules labeled with Alexa Fluor 647. This plane intersects the lateral plane (inset) along the white dashed line. (**b**) Direct 45° obSTORM image of **a**. Inset: a frame image of single molecules for the green dashed region. (**c, d**) Distributions of the number of photons detected per molecule in one frame, *N*, and localization precision, σ_*x*_ and σ_*o*_, in the *x* and *o* directions. (**e**) Zoomed view of the white boxed area in **b** and the intensity profile (blue) across the two microtubules (after binned along the length of the microtubules within the white dashed area) compared with the profile (gray) for the same region from **a**. (**f**) Diffraction-limited, axial plane (α = 90°) fluorescence image of microtubules of an A549 cell cluster labeled with CF568. (**g**) Direct 90° obSTORM image of **f**. (**h, i**) Distributions of signal photons and localization precision. (**j**) Magnified views of the two boxed regions in **g** and the cross-sectional profiles (after binned as in **e**). (**k**) Two-color 45° obSTORM image of microtubules (magenta, Alexa Fluor 647) and mitochondria (green, CF568). The inset shows the conventional fluorescence, lateral view image with the intersection line. (**l, m**) Zoomed views of the two boxed regions in **k**. Scale bars, 2 μm (**b**, inset in **b, g, k**), 10 μm (insets in **a, k**), 200 nm (**e, j**) and 500 nm (**l, m**).

The average localization precision^18^ in standard deviation was isotropically ∼18.5 nm in the oblique plane throughout the depth of the sample (**Fig. 2d**), corresponding to an imaging resolution of ∼44 nm in FWHM. Comparable results were consistently found over different cells (**Supplementary Fig. 7**). These values compare favorably to 3D-STORM^4^, which typically gives 25-30 nm lateral resolution and ∼60 nm axial resolution for a depth range of ∼800 nm within a few microns from the coverslip, but the resolution quickly deteriorates deeper into the sample. Two adjacent tubules were clearly resolved at a 128 nm center-to-center distance (**Fig. 2e**), in contrast to their diffraction-limited fluorescence image. The apparent widths of both microtubules were ∼69 nm, consistent with a convolution of our resolution with the width of antibody-labeled microtubules (56 nm)^19^. A network of mitochondria in A549 cells was also successfully imaged in 45° obSTORM with a clear visualization of mitochondrion’s hollow shapes (**Supplementary Fig. 8**).

We further demonstrate 90° obSTORM in which the axial plane is directly imaged (**Fig. 2f,g**). The increased pupil loss at 90° further lowers the relative collection efficiency down to ∼10%, resulting in the average photon count from CF568 probes of ∼500 (**Fig. 2h**). The elliptical PSF at 90° leads to anisotropic localization precisions that averaged out to 22 nm and 48 nm in the *x* and *z* directions, respectively (**Fig. 2i**), corresponding to FWHM resolutions of 52 nm (*x*) and 114 nm (*z*). In **Fig. 2j**, the measured width of microtubules was 77 nm (*x*) and 126 nm (*z*), both of which are again consistent with a convolution of our resolution with the anticipated width of antibody-labeled microtubules (56 nm). The reduced resolution in 90° obSTORM still resolved the hollow structure of mitochondria (**Supplementary Fig. 9**). The 90° obSTORM resolution may potentially be enhanced to <50 nm with further improvement (**Supplementary Discussion**).

We next demonstrate three additional capabilities of obSTORM. First, obSTORM is fully compatible with multi-color super-resolution imaging. For example, in our two-color obSTORM images, mitochondria and microtubules in A549 cells are both well resolved, and the former were correctly surrounded by the latter (**Fig. 2k-m** and **Supplementary Fig. 10**). Comparable results were obtained with opposite combinations of AF647 and CF568 labeling of the two targets (**Supplementary Fig. 11**), and we obtained good results for both 45° and 90° oblique angles (**Supplementary Fig. 12**). Second, obSTORM enables two-plane super-resolution imaging where both an oblique plane and a lateral plane are captured simultaneously by two cameras (**Supplementary Fig. 10-12**). With 50% signal loss at the PBS, the lateral-plane STORM yielded an average localization precision of 10-15 nm. Third, another dimension of localization normal to the oblique plane can be added in obSTORM. We calibrated the astigmatic PSF of the obSTORM system augmented by a cylindrical lens for 3D localization and reconstructed a 3D, 45° obSTORM image of mitochondria in an A549 cell with a section thickness of 1.5 μm (**Supplementary Fig. 13**).

obSTORM achieves uniform super-resolution throughout depth in cell and tissue samples. For over-confluence A549 cells, we obtained a 45° obSTORM image of microtubule structures extending over 16.3 μm in the *o* direction, equivalent to 11.5 μm in the conventional *z* direction (**Fig. 3a**). Two microtubules close to and far away from the coverslip (*z* = 0.9 μm and 9.2 μm, respectively) were imaged similarly in diameter (**Fig. 3b**). More rigorously, the average number of collected photons and the average localization precision evaluated through imaging depth indicated uniform resolution (**Fig. 3c**). A colony of A549 cells that extended deeper up to *z* = 14.4 μm was also imaged uniformly (**Supplementary Fig. 8**). For tissue samples, we imaged the AF647-labeled synaptonemal complex (SYP-4) of *Caenorhabditis elegans* gonads immobilized in a 1% agarose gel between a coverslip and a glass slide. A 45° cross-sectional STORM image of *C. elegans* was obtained up to *z* = 32 μm in depth from the coverslip surface (**Fig. 3d**). Image features ∼87 nm in width were well-resolved at ∼125 nm center-to-center distances (**Fig. 3e**). The through-depth analysis of localization data revealed good uniformity over the 32 μm depth in single-molecule photon count and localization precision (**Fig. 3f**), with a near-isotropic imaging resolution of ∼51 nm in FWHM.

**Figure 3.**
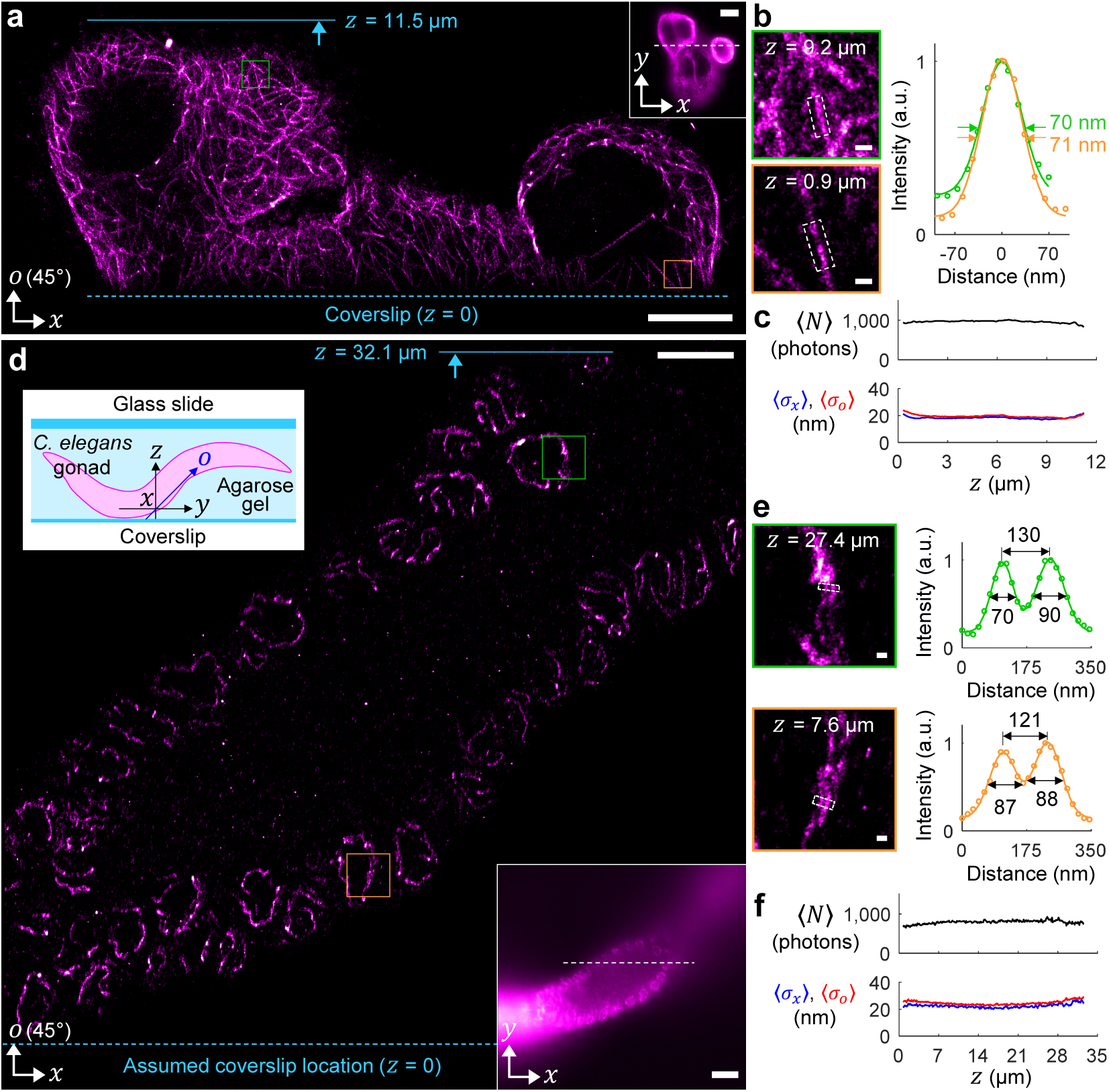
45° obSTORM with thick samples. (**a**) Super-resolution 45° oblique plane image of microtubule structure labeled by Alexa Fluor 647 in A549 cells. The inset shows the conventional fluorescence, lateral image with the intersection line. (**b**) Zoomed views of the two boxed regions at different depths and the intensity profiles of individual microtubules after averaged along their direction over the dashed box areas. (**c**) Through-depth distributions of the average number of signal photons, ⟨*N*⟩, and the average localization precisions, ⟨σ_*x*_⟩ and ⟨σ_*o*_⟩. (**d**) 45° obSTORM image of SYP-4 in *C. elegans* germ cells labeled by Alexa Fluor 647. Two insets show the illustration of a worm gonad with the oblique plane where the image is taken (upper left) and the conventional fluorescence, lateral image with the intersection of two image planes (lower right). (**e**) Magnified views of the synaptonemal complex at 7.6 μm and 27.4 μm depths and their cross-sectional profiles. (**f**) Distributions of ⟨*N*⟩, ⟨σ_*x*_⟩ and ⟨σ_*o*_⟩ through sample depth. Scale bars, 5 μm (**a, d**), 10 μm (insets in **a, d**), 200 nm (**b, e**).

We demonstrated a new technique, obSTORM, that offers single-shot super-resolution imaging for any oblique plane through deep inside thick samples. A uniform, isotropic resolution of 44-51 nm was achieved in 45° oblique view throughout a depth up to 32 μm in cell and tissue samples, and we showed with fluorescent beads that comparable resolutions may be achieved for imaging depths of ∼100 μm (**Supplementary Fig. 5**). With further improvement, the resolution of obSTORM may reach <50 nm along any directional view (**Supplementary Discussion**). This method can capture multi-color, large 3D volumetric super-resolution images from oblique stacks and should be compatible with live-cell samples as well as other probes such as fluorescent proteins and quantum dots. obSTORM thus opens a new door for direct, facile cross-sectional super-resolution imaging of cells, tissues and potentially whole animals.

## Author contributions

J.K. designed and built the microscope system, wrote software codes for hardware control and localization analysis, and conducted theoretical calculations. J.K. prepared fluorescent bead samples and calibrated the system. M.W. prepared cell and tissue samples. J.K. and M.W. carried out the imaging experiments. J.K. and Y.W. analyzed the data. X.Z. and K.X. guided the research. All authors contributed to discussions and writing of the manuscript.

## Acknowledgments

We thank L. Li for providing gold-evaporated silicon wafer mirrors and S. Köhler (Dernburg Lab) for helping with *C. elegans* samples. This work was supported by Gordon and Betty Moore Foundation. K.X. is a Chan Zuckerberg Biohub investigator and acknowledges support from the Bakar Fellows Award. M.W. acknowledges NSF Graduate Research Fellowship (DGE-1106400).

## METHODS

### The optical setup of obSTORM

The setup was built on an optical table as illustrated in **Supplementary Fig. 1**. For illumination, three diode-pumped solid-state (DPSS) lasers were used for excitation at 641 nm (40 mW, OBIS 640LX-40, Coherent) and 532 nm (1300 mW, Newgazer Technology, China) and for activation at 405 nm (300 mW, Newgazer Technology, China). Each laser beam was individually coupled to single mode fibers (SMF, S405-XP, Thorlabs), collimated to obtain a quality beam profile with the e^−2^ diameters of 1.3-1.6 mm, and then combined by two dichroic beam splitters (T588lpxr-UF2, Chroma and Di03-R488-t1-25×36, Semrock). The circular beams were shaped to slit beams by cylindrical focusing (CL, *f* = 250 mm, ACY254-250-A, Thorlabs) and cylindrical beam expansion with a pair of achromatic cylinder lenses (*f* = 50 mm, ACY254-050-A, Thorlabs and *f* = 200 mm, ACY254-200-A, Thorlabs). The slit beam formed on a gimbal mirror for beam steering was demagnified by a 4f system, comprising the L1 lens (*f* = 200 mm, ACH254-200-A-ML, Thorlabs) and a water immersion objective (O1, UPLSAPO 60XW/1.20, Olympus) mounted to a PZT scanner (P-721, Physik Instrumente), with a dichroic beam splitter (DBS, Di03-R405/488/532/635-t1, Semrock). The e^−2^ thickness of in-focus light-sheets at α = 90° was ∼1.7 and ∼2.1 µm for the 532 nm and 641 nm light, respectively. Two mirrors with flipper optic mounts (9891, Newport) allowed a bypass for wide-field illumination through a 5.4× beam expansion with two lenses (M-10X, Newport and #49-782, Edmund Optics). The oblique light-sheet illumination covers α = 45-90° and the wide-field illumination can cover α = 0-45° at compromised signal-to-background ratio. Samples mounted between standard microscope coverslips and slides (75 mm × 25 mm) were loaded to a home-made sample holder on a 3D stage (562-XYZ, Newport).

For fluorescence detection, the O1 was relayed with a remote objective (O2, MPlanApo 50x/0.95, Olympus) via a 4f system with two achromatic lenses (*f* = 200 mm, #47-645, Edmund Optics and *f* = 180 mm, AC508-180-A-ML, Thorlabs). Three remote mirrors (M) were prepared: a protected silver mirror (PF10-03-P01, Thorlabs) for α = 0°, a home-made gold-evaporated Si wafer mirror (10 mm × 15 mm) fixed on the 22.5° surface of a pentaprism (PS-932, Thorlabs) for α = 45°, and a knife-edge right-angle prism mirror (MRAK25-P01, Thorlabs) for α = 90°. These mirrors were housed by or glued on lens tubes (SM1Lxx, Thorlabs) and easily switched at a kinematic rotation mount (KS1RS, Thorlabs) on a 2D stage (462-XY-SD, Newport) driven by PZT micrometers (NPM140SG, Newport). In principle, a single small mirror can cover all oblique angles (0-90°) with its rotation/translation controlled. A polarizing beam splitter cube (CCM1-PBS251, Thorlabs) and an achromatic quarter-wave plate (AQWP05M-600, Thorlabs) direct the remote fluorescence signals to an electron-multiplying charge-coupled device (EMCCD) camera (EMCCD1, Cascade II 512, Photometrics) via a 4f system of the O2 and the L4 lens (*f* = 300 mm, AC254-300-A-ML). A simultaneous lateral (*xy*) image is formed by the L5 lens (*f* = 200 mm, AC254-200-A-ML, Thorlabs) on the EMCCD2 camera (Luca R, Andor Technology). Quad-band emission filters (FF01-446/510/581/703-25, Semrock) were used for AF647 and a bandpass filter (FF01-590/104-25, Semrock) was added for CF568.

### System alignment and calibration

To accurately construct all 4f configurations, we measured the front focal length and back focal length of all achromatic doublets and the location of back focal plane of objectives used. In our optical design and alignment, we considered the finite thickness of DBS, PBS, and QWP through which uncollimated fluorescence light passes.

The system’s 3D magnification in the remote focusing^20^ was identified for localization data correction. We imaged a group of isolated fluorescent beads with EMCCD2 at a reference focus of O1 (*z* = 0) and then at α = 0° with EMCCD1 at the remote focus of O2 (*z*_*r*_ = 0) (**Supplementary Fig. 6**). The O1 was 10 μm-stepped across 100 μm and the corresponding remote focus was found with brightest bead images on EMCCD1. From this series of bead images, we extracted a change of lateral magnification over depth. We set a compensator of the L3-O2 distance to uniformize the magnification at 685 nm emission (<0.5% distortion, **Supplementary Fig. 6**). The absolute magnification (or image pixel size) at focus (*z* = 0) was also calibrated with a microscope calibration slide (MR400, AmScope).

For two-color obSTORM imaging, we quantified the chromatic focal shift in the detection system (sample to EMCCD1) using a multi-color fluorescent bead (T7280, Invitrogen) as detailed in **Supplementary Fig. 10**. After the amount of focal shift in the orange channel (570 nm) relative to the dark-red channel (685 nm) was found, each channel’s in-focus PSF images was localized to extract the pixel registration error. This calibration was performed at α = 45° and 90° separately and applied during STORM data acquisition and image reconstruction. We also fine-tuned the illumination optics that combines the three laser beams to accurately overlap the light-sheets when seen on EMCCD1 after considering the chromatic focal shift present in the fluorescence detection optics.

### Sample preparation

#### Fluorescent bead samples

For system characterization and calibration purposes, we used several fluorescent beads (F8800, T7284, P7220, and T7280, Invitrogen). The bead solution was dried on plasma-etched #1.5H coverslips directly or after diluted in deionized (DI) water to keep good separation. The bead samples were mounted with DI water on glass slides.

#### Immunofluorescence of A549 cells

Cells were cultured following standard tissue culture protocols and plated on #1.5H coverslips (CG15CH, Thorlabs) at ∼60% confluency. After 24 h, cells were fixed using a solution of 3% paraformaldehyde and 0.1% glutaraldehyde in phosphate-buffered saline followed by two washes with 0.1% sodium borohydride in phosphate-buffered saline. Cells were subsequently blocked and permeabilized in a buffer containing 3% bovine serum albumin with 0.5% Triton X-100, and incubated overnight at 4°C in primary antibody solutions with anti-alpha tubulin (Mouse monoclonal, Clone DM1α, 1:500, T9026, Sigma) to label microtubules, and anti-Tom20 (Rabbit polyclonal, 1:200, sc-11415, Santa Cruz Biotechnology) to label the outer membranes of mitochondria. After triple-washing, cells were incubated for 45 min at room temperature in secondary antibody solution with AF647-anti-Mouse (1:500, A31571, Invitrogen) or AF647-anti-Rabbit (1:500, A21245, Invitrogen) and CF568-anti-Mouse (1:100, 20827, Biotium) or CF568-anti-Rabbit (1:100, 20828, Biotium). Stained cells were mounted in ∼10 μL STORM imaging buffer (Tris-HCl, pH 7.5, containing 100 mM cysteamine, 5% glucose, 0.8 mg/mL glucose oxidase, and 40 μg/mL catalase) and sealed prior to imaging.

#### Immunofluorescence of C. elegans

Dissected gonads from young adults 24 h post-L4 were immunostained as previously described in literature^21^, ^22^. Fixed tissue was placed in small polypropylene tubes and stained in suspension. The primary antibody was anti-HA (mouse monoclonal 2–2.2.14, 1:500, ThermoFisher Scientific) used on an SYP-4 HA knock-in mutant created by using CRISPR-Cas9 as described^22^ and the secondary antibody was AF647-anti-Mouse (1:500, A31571, Invitrogen). Stained tissue in polypropylene tubes was then immersed in 100 μL imaging buffer (Tris-HCl, pH 7.5, containing 100 mM cysteamine, 5% glucose, 0.8 mg/mL glucose oxidase, and 40 μg/mL catalase), mixed with 100 μL agarose solution (2%, 60°C), and immediately transferred to a heated microscope slide on a hot plate at 50°C. Four-sided Kapton tape spacers were used on the glass slide. The sample was covered with a pre-heated #1.5H coverslip at 50°C, quickly cooled down to room temperature to form ∼1% agarose gel for sample immobilization, and sealed by nail polish.

### Image acquisition

A prepared sample slide was mounted on the sample holder, and water immersion oil (Immersol W 2010, 444969, Carl Zeiss) was applied. We first located samples in conventional fluorescence imaging with EMCCD2 under wide-field illumination with a peak power density of 5-20 W/cm^2^. Then an oblique plane fluorescence image was checked on EMCCD1 under weak light-sheet illumination (20-80 W/cm^2^). Prior to STORM image acquisition, samples were excited at the highest laser power for 0-3 minutes to turn off most fluorophores and reduce background fluorescence. Then video data was recorded at 28-50 frames/s (exposure time: 20-35 ms) until 20,000-60,000 frames were acquired per each color channel. The dye channel with longer emission wavelength (AF647) was imaged first, followed by the CF568 channel at a corrected, chromatic object focus to obtain the STORM data on the identical plane. A typical light-sheet power was 10-25 kW/cm^2^ and 30-70 kW/cm^2^ at 641 nm and 532 nm light, respectively. A good single-molecule density was maintained during image acquisition by increasing weak 405 nm activation light within a range of 0-10 W/cm^2^.

### Single-molecule localization and image reconstruction

Two-dimensional localization of single-molecule images was performed as follows. After locating each molecule’s position at pixel accuracy from bandpass filtered images as detailed at http://site.physics.georgetown.edu/matlab, we fitted raw single-molecule images at a fixed window size with a 2D elliptical Gaussian function plus a constant background, 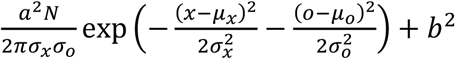, where *a* denotes the pixel size, *N* the number of collected photons, *b*^*^2^*^ the number of background photons per pixel, and *µ*_*i*_ and *σ*_*i*_ the center position and the pixelated size of Gaussian PSF along the i axis (*i* = *x, o*), respectively^18^. Using this fitting result as an initial guess with an updated window size of ∼2×FWHM along each axis, each molecule image was fitted again by the least square (LS) algorithm solved by MATLAB’s “lsqcurvefit” function. The resultant localization precision^18^ could be derived as 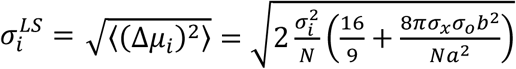 where an excess noise factor of ∼2 in EMCCD cameras^23^ was assumed. Then, the whole localization data was compensated as follows. The non-uniform image magnification was first corrected (**Supplementary Fig. 6**). In case of two-color data, the relative pixel registration of the orange color channel was adjusted (**Supplementary Fig. 10**). The drift of localization data during data acquisition was estimated and cancelled out by the correlation method implemented in PALMsiever^24^. The same localization procedure was applied for lateral-plane STORM data if acquired simultaneously.

The final localization data was binned (10-15 nm grid) and smoothed with a 2D Gaussian filter (sigma: 5-15 nm) by in-house MATLAB reconstruction codes.

## References

1. Rust, M.J., Bates, M. & Zhuang, X. Sub-diffraction-limit imaging by stochastic optical reconstruction microscopy (STORM). Nature Methods 3, 793–795 (2006).

2. Betzig, E. et al. Imaging intracellular fluorescent proteins at nanometer resolution. Science 313, 1642–1645 (2006).

3. Hess, S.T., Girirajan, T.P.K. & Mason, M.D. Ultra-High Resolution Imaging by Fluorescence Photoactivation Localization Microscopy. Biophysical Journal 91, 4258–4272 (2006).

4. Huang, B., Wang, W., Bates, M. & Zhuang, X. Three-dimensional super-resolution imaging by stochastic optical reconstruction microscopy. Science 319, 810–813 (2008).

5. Pavani, S.R.P. et al. Three-dimensional, single-molecule fluorescence imaging beyond the diffraction limit by using a double-helix point spread function. Proc. Natl. Acad. Sci. U. S. A. 106, 2995–2999 (2009).

6. Shechtman, Y., Sahl, S.J., Backer, A.S. & Moerner, W.E. Optimal Point Spread Function Design for 3D Imaging. Physical Review Letters 113, 133902 (2014).

7. Huang, B., Jones, S.A., Brandenburg, B. & Zhuang, X.W. Whole-cell 3D STORM reveals interactions between cellular structures with nanometer-scale resolution. Nat. Methods 5, 1047–1052 (2008).

8. Tokunaga, M., Imamoto, N. & Sakata-Sogawa, K. Highly inclined thin illumination enables clear single-molecule imaging in cells. Nat Meth 5, 159–161 (2008).

9. Cella Zanacchi, F. et al. Live-cell 3D super-resolution imaging in thick biological samples. Nat Meth 8, 1047–1049 (2011).

10. Gebhardt, J.C.M. et al. Single-molecule imaging of transcription factor binding to DNA in live mammalian cells. Nat Meth 10, 421–426 (2013).

11. Galland, R. et al. 3D high-and super-resolution imaging using single-objective SPIM. Nat Meth 12, 641–644 (2015).

12. Meddens, M.B.M. et al. Single objective light-sheet microscopy for high-speed whole-cell 3D super-resolution. Biomedical Optics Express 7, 2219–2236 (2016).

13. Dunsby, C. Optically sectioned imaging by oblique plane microscopy. Opt Express 16, 20306–20316 (2008).

14. Kumar, S. et al. High-speed 2D and 3D fluorescence microscopy of cardiac myocytes. Opt Express 19, 13839–13847 (2011).

15. Anselmi, F., Ventalon, C., Begue, A., Ogden, D. & Emiliani, V. Three-dimensional imaging and photostimulation by remote-focusing and holographic light patterning. Proc. Natl. Acad. Sci. USA 108, 19504–19509 (2011).

16. Kim, J., Li, T., Wang, Y. & Zhang, X. Vectorial point spread function and optical transfer function in oblique plane imaging. Opt Express 22, 11140–11151 (2014).

17. Li, T. et al. Axial Plane Optical Microscopy. Scientific Reports 4, 1–6 (2014).

18. Mortensen, K.I., Churchman, L.S., Spudich, J.A. & Flyvbjerg, H. Optimized localization analysis for single-molecule tracking and super-resolution microscopy. Nat Meth 7, 377–381 (2010).

19. Bates, M., Huang, B., Dempsey, G.T. & Zhuang, X. Multicolor Super-Resolution Imaging with Photo-Switchable Fluorescent Probes. Science 317, 1749 (2007).

20. Botcherby, E.J., Juskaitis, R., Booth, M.J. & Wilson, T. Aberration-free optical refocusing in high numerical aperture microscopy. Opt Lett 32, 2007–2009 (2007).

21. Phillips, C.M., McDonald, K.L. & Dernburg, A.F. in Meiosis: Volume 2, Cytological Methods. (ed. S. Keeney) 171–195 (Humana Press, Totowa, NJ; 2009).

22. Köhler, S., Wojcik, M., Xu, K. & Dernburg, A.F. Superresolution microscopy reveals the three-dimensional organization of meiotic chromosome axes in intact Caenorhabditis elegans tissue. Proceedings of the National Academy of Sciences 114, E4734–E4743 (2017).

23. Hynecek, J. & Nishiwaki, T. Excess noise and other important characteristics of low light level imaging using charge multiplying CCDs. IEEE Transactions on Electron Devices 50, 239–245 (2003).

24. Pengo, T., Holden, S.J. & Manley, S. PALMsiever: a tool to turn raw data into results for single-molecule localization microscopy. Bioinformatics 31, 797–798 (2015).

