## Supplementary Materials for "Oblique plane single-molecule localization microscopy for thick samples"



| | |
|---|---|
| **Supplementary Figure 1** | The complete layout of obSTORM |
| **Supplementary Figure 2** | Effect of polarization optical components on PSF |
| **Supplementary Figure 3** | Theoretical PSF at potentially higher NA configurations |
| **Supplementary Figure 4** | Theoretical and experimental PSF at $\lambda$ = 570 nm |
| **Supplementary Figure 5** | An imaging depth of obSTORM estimated by fluorescent beads |
| **Supplementary Figure 6** | Calibration of 3D imaging magnification |
| **Supplementary Figure 7** | Other 45° obSTORM images of microtubules in A549 cells |
| **Supplementary Figure 8** | 45° obSTORM images of mitochondria in A549 cells |
| **Supplementary Figure 9** | 90° obSTORM images of mitochondria in A549 cells |
| **Supplementary Figure 10** | Calibration for two-color 45° obSTORM and another example of two-color 45° obSTORM image of an A549 cell |
| **Supplementary Figure 11** | Two other examples of two-color 45° obSTORM images with dye molecules switched in A549 cells |
| **Supplementary Figure 12** | Two-color 90° obSTORM images of microtubules and mitochondria in A549 cells |
| **Supplementary Figure 13** | 3D super-resolution imaging in 45° obSTORM |
| **Supplementary Derivation** | Theoretical PSF in obSTORM |
| **Supplementary Discussion** | Potential improvement on resolution in obSTORM |


Note: **Supplementary Video 1** is available online.

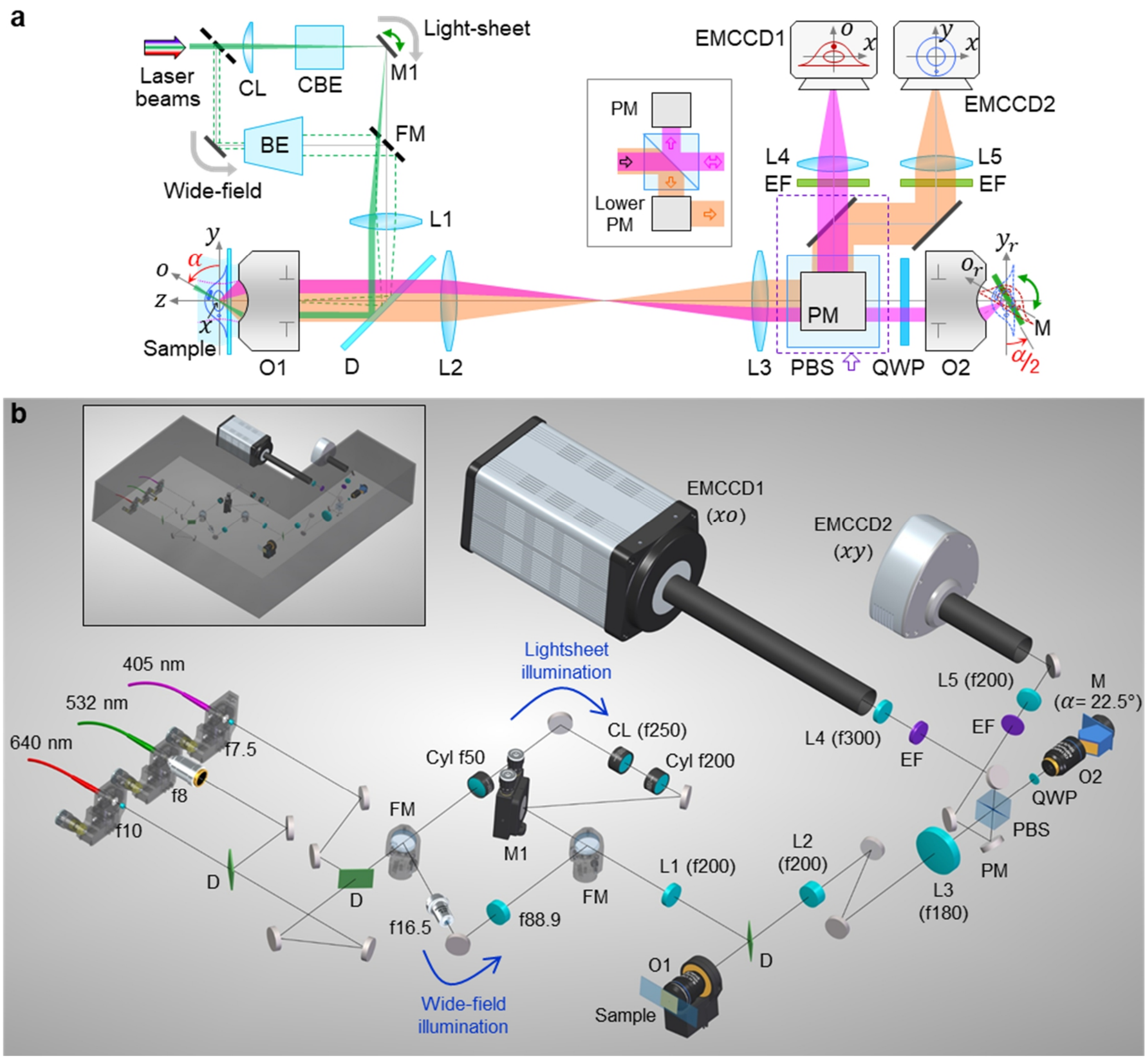


**Supplementary Figure 1 | The complete layout of obSTORM.** (**a**) Top view of the system layout. The illumination can be wide-field or light-sheet (green paths). A slit beam, formed by a cylinder lens (CL) and a cylindrical beam expander (CBE), is steered by a mirror (M1) and demagnified in sample space to generate an oblique light-sheet with a tunable $\alpha$ of 45-90°. An oblique fluorescence image along the excited $xo$ plane is refocused by remote objective (O2), transformed to the focal plane ($x_r y_r$) by a tilted mirror (M), and then directly captured by EMCCD1 via the periscope mirror (PM) above PBS (magenta path). A lateral ($xy$) fluorescence image captured from sample objective (O1) can be simultaneously obtained by EMCCD2 via another periscope underneath PBS (orange path). The inset shows the front view of the area enclosed with the dashed violet box. FM, flip mirror; EF, emission filter. (**b**) The actual 3D experimental layout. The focal length of each lens used (in millimeters) is labeled. Beam expanders are used in wide-field beam path by two lenses (f16.5 and f88.9) and in light-sheet beam path by two cylindrical lenses (f50 and f200). The inset shows an optical enclosure (excluding the laser heads and EMCCD cameras) to improve mechanical/thermal stability and to avoid unwanted room light.

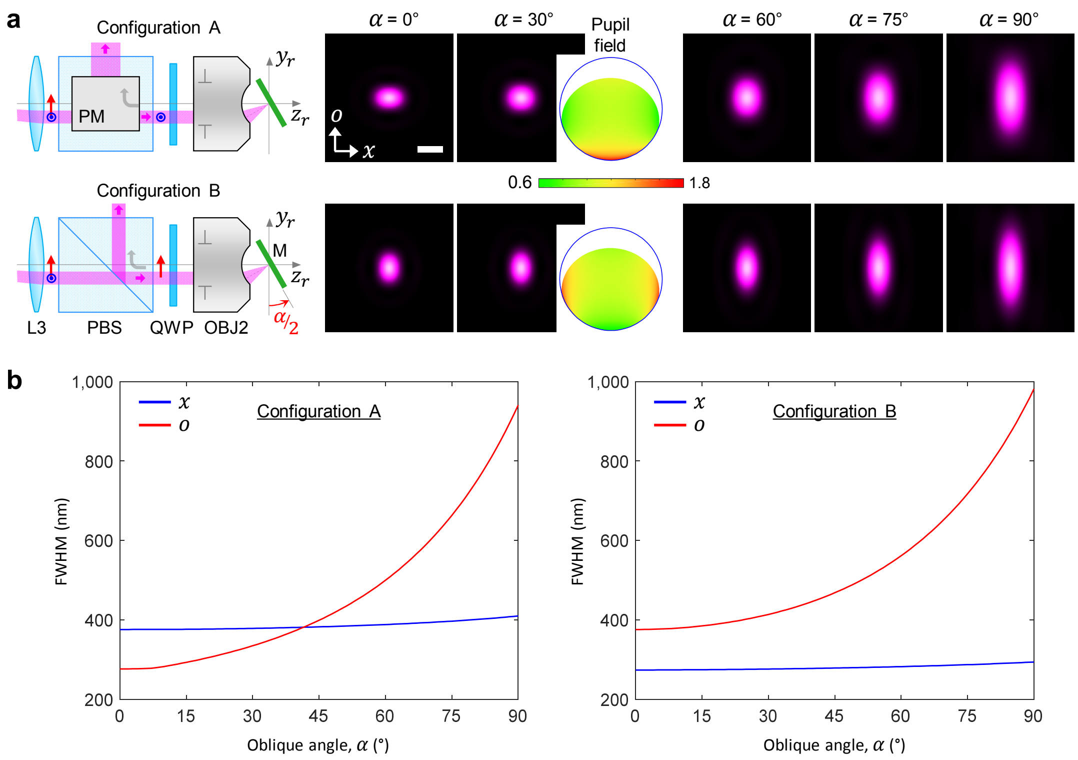


**Supplementary Figure 2 | Effect of polarization optical components on PSF.** (**a**) In Configuration A, only $p$ wave (a.k.a. TM wave, shown as blue concentric circles) component of single-molecule fluorescence transmits through the polarizing beam splitter (PBS) for oblique plane imaging. The transmitted wave is primarily from the dipole emitter orientated along $x$ and yields stronger electric field strength at the edge of the exit pupil in the $o$ direction as shown in the case of $\alpha = 30°$. As PSF is conceptually a Fourier transform of pupil field distribution, such strong edge field helps to compensate the pupil loss occurring in the same $o$ direction and thus slow down the PSF elongation over oblique angle. This means that the reshaped pupil field distribution by the PBS can favorably balance the intrinsically asymmetric pupil loss[1] in oblique plane microscopy and therefore less elliptical and overall smaller PSF is achievable. Such a balance is maximized at the highest semi-aperture angle of objectives (**Supplementary Fig. 3**). On the other hand, in Configuration B (found in a literature[2]), $s$ wave (or TE wave, indicated as red arrows) is instead used, which produces rather weaker field strengths in the $o$ direction and thus yields even more anisotropic PSF that is vertically elongated over all oblique angles. This trend is clearly shown in theoretical 2D PSFs calculated at $\lambda = 685$ nm ($NA_1 = 1.2$ (water immersion), $NA_2 = 0.95$) based on **Supplementary Derivation**. Our experimental setup uses the Configuration A. Scale bar, 400 nm. (**b**) Theoretical FWHM of PSF in **a** for each configuration. The Configuration A predicts a circular PSF of 382 nm in FWHM at $\alpha = 41.5°$, while the Configuration B predicts an elliptical PSF with an undesirably larger FWHM of 453 nm in the $o$ direction. The use of PBS and QWP doubles light efficiency compared with that of a non-polarizing beam splitter (NPBS) alone (although not shown here). The PSF in the latter case is circular only at $\alpha = 0°$ and starts to stretch vertically as the oblique angle increases[1].

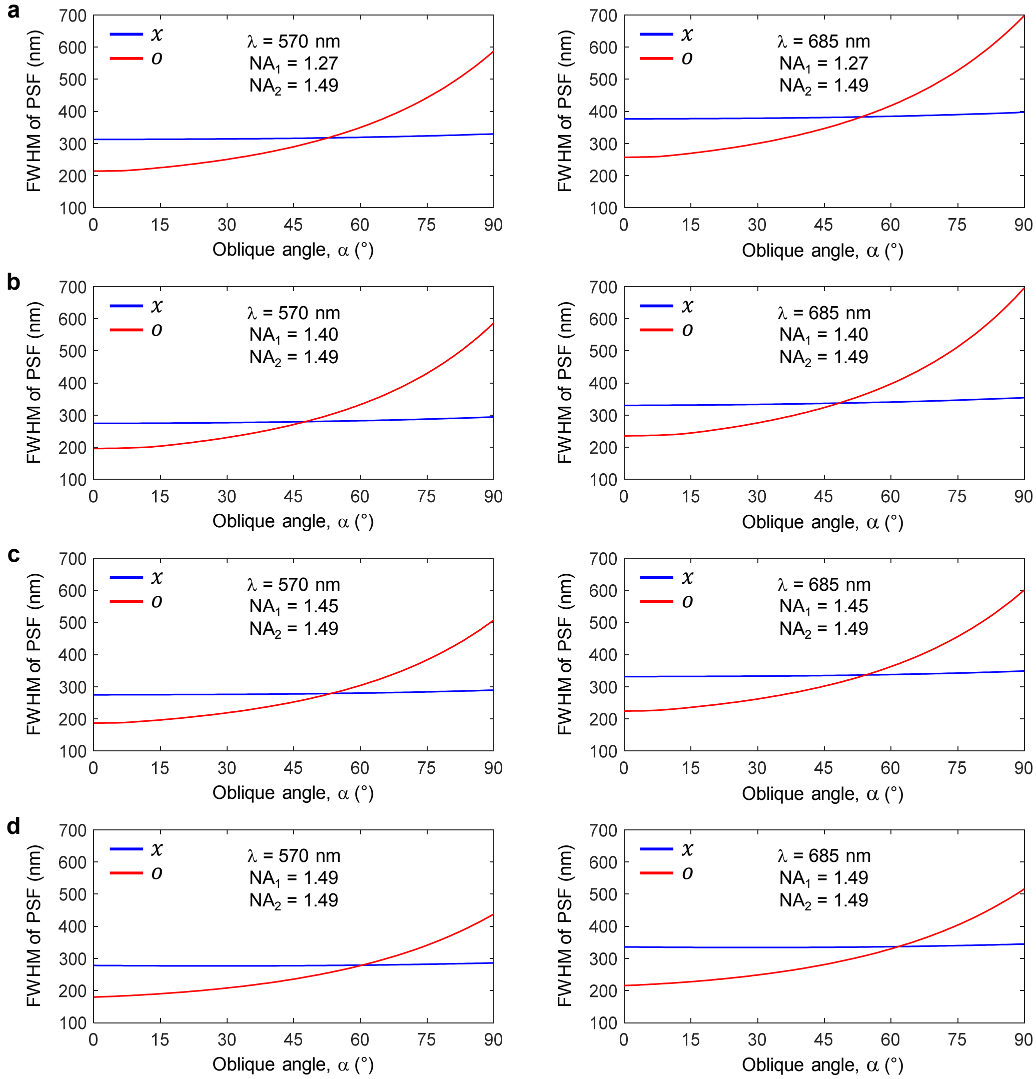


**Supplementary Figure 3 | Theoretical PSF at potentially higher NA configurations.** $NA_1$, NA of the sample objective (O1); $NA_2$, NA of the remote objective (O2). FWHMs of PSFs with 1.27 NA/1.49 NA (**a**), 1.4 NA/1.49 NA (**b**), 1.45 NA/1.49 NA (**c**) and 1.49 NA/1.49 NA (**d**) at emission of 570 nm (left column) and 685 nm (right column). The assumed immersion is water (refractive index $n$ = 1.333 at 570 nm and 1.330 at 685 nm) in **a** and oil ($n$ = 1.516 at 570 nm and 1.512 at 685 nm) in **b**-**d**. Higher cone-angle systems predict isotropic PSFs at higher oblique angles at 53°, 48°, 54° and 61° in **a**, **b**, **c** and **d**, respectively. The 1.49 NA/1.49 NA system gives least elliptical and smallest PSF over all oblique angles, which may benefit SMLM. Compared with the 1.2-NA/0.95-NA system in **Fig. 1c** at α = 90°, the FWHM in the *o* direction at λ = 685 nm is reduced to 698/696/601/516 nm in **a**/**b**/**c**/**d** with the increased pupil areas by 1.63×/1.43×/1.67×/1.96× calculated based on **Supplementary Derivation**. Thus, these higher NA systems can further improve the localization precision approximately by 1.7×, 1.6×, 2.0×, and 2.6× in **a**, **b**, **c**, and **d**, respectively in terms of $FWHM/\sqrt{N}$, where $N$ denotes the number of collected photons which is roughly proportional to the pupil area here.

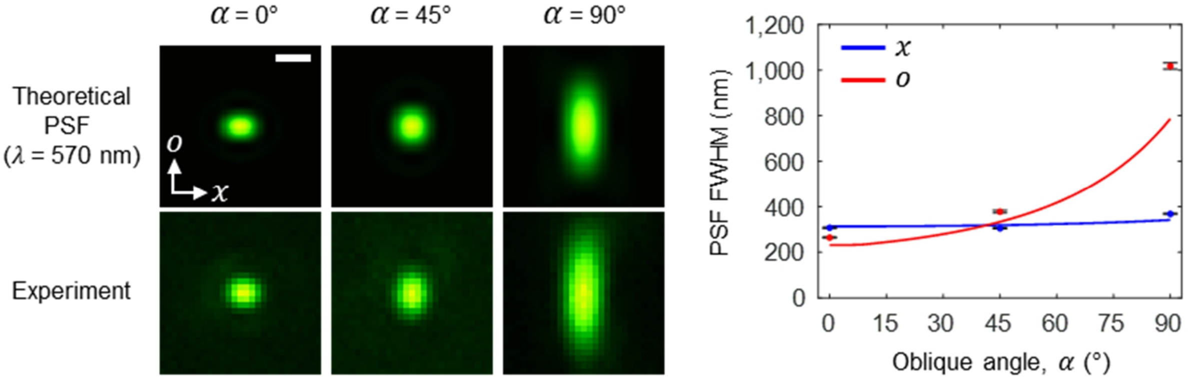


**Supplementary Figure 4 | Theoretical and experimental PSF at λ = 570 nm.** Comparison of theoretical PSFs and experimental PSFs from orange fluorescent beads (F8800, Invitrogen for α = 0° and 45°, and P7220, Invitrogen for α = 90°). On the right, the theoretical (line) and experimental (dots with error bars) FWHMs in the *x* and *o* directions over oblique angles show a similar trend shown in **Fig. 1b,c** at 685 nm. The water-immersion, objective NA in the sample space is 1.2 (refractive index: 1.333 at 570 nm), and the remote NA is 0.95. PSFs in **Fig. 1b,c** and here were captured with an 8-µm-pixel EMCCD camera (not by the 16-µm-pixel EMCCD1) for finer spatial sampling. Error bars, s.d.; *n* = 10, 6, and 8 beads for 0, 45, and 90° respectively. Scale bar, 400 nm.

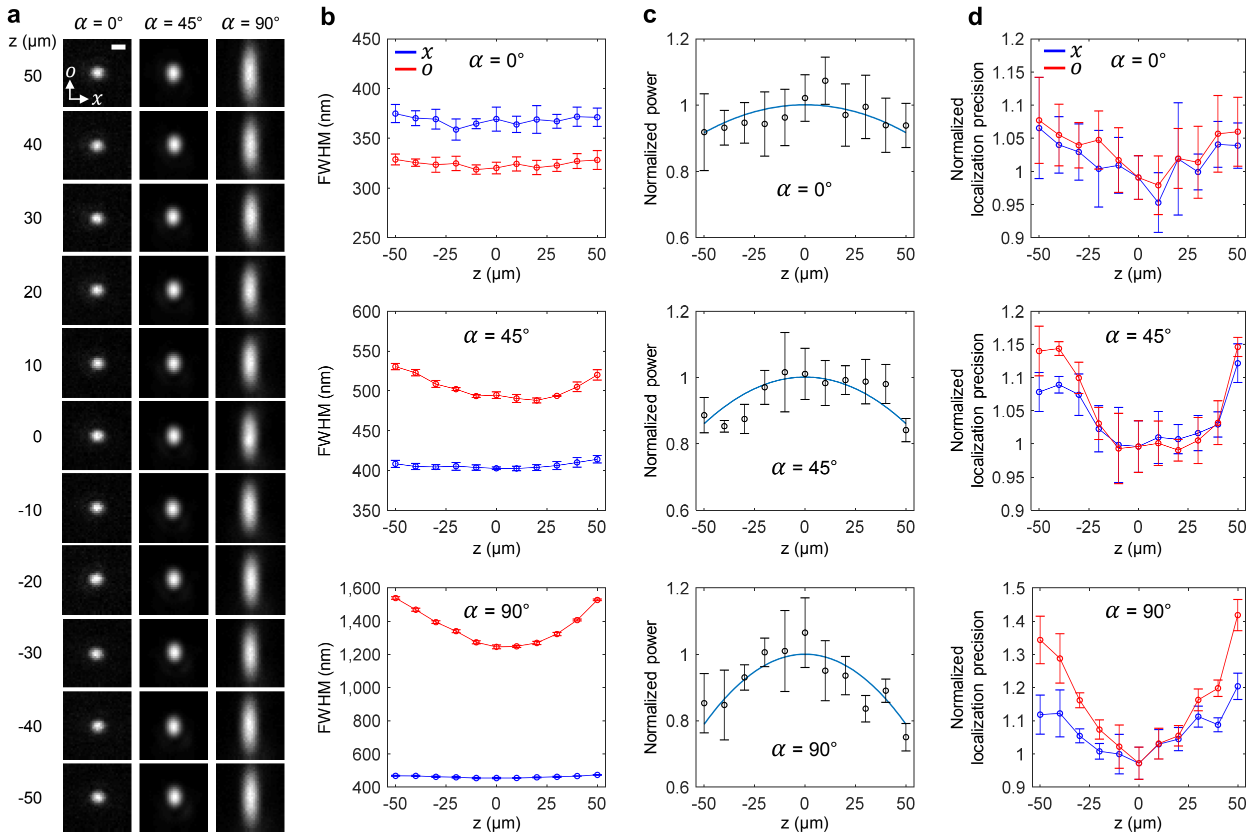


**Supplementary Figure 5 | An imaging depth of obSTORM estimated by fluorescent beads.** (**a**) Measured PSFs across 100 μm in axial depth. The sample objective (O1) was stepped by 10 μm with respect to a dark-red fluorescent bead sample (⌀100 nm, T7284, Invitrogen for 0° and ⌀175 nm, P7220, Invitrogen for 45° and 90°) and then the focal position of the remote tilting mirror (M) was adjusted accordingly. The PSF images were taken with an 8-μm-pixel EMCCD camera (not with the 16-μm-pixel EMCCD1) for finer spatial sampling. Scale bar, 500 nm. (**b**) The FWHMs of the measured PSFs at three oblique angles. (**c**) The normalized beam power of PSF over the 100 μm depth and the quadratic fit (sky blue line). At 0° oblique angle, through-focus PSF is uniform with less than 10% power loss over the 100 μm depth, implying negligible optical aberrations present in the aligned system. At α = 45° and 90°, the modest increase in the PSF size together with quadratic power loss through focus was observed. (**d**) The relative localization precision through depth estimated simply by $FWHM/\sqrt{Power}$ from **b** and **c**. The localization precision deteriorates <15% (<40%) at the oblique angle of 45° (90°) for the 100 μm depth. An axial range of <15% deterioration in localization precision at 90° oblique plane is ~60 μm. This through-depth estimation with fluorescent beads does not consider the scattering effect of samples in finite thickness, which is typically negligible over a depth of ~100 μm in many weakly-scattering samples[3]. Error bars, s.d., $n$ = 9, 4, and 6 beads for 0°, 45°, and 90°, respectively.

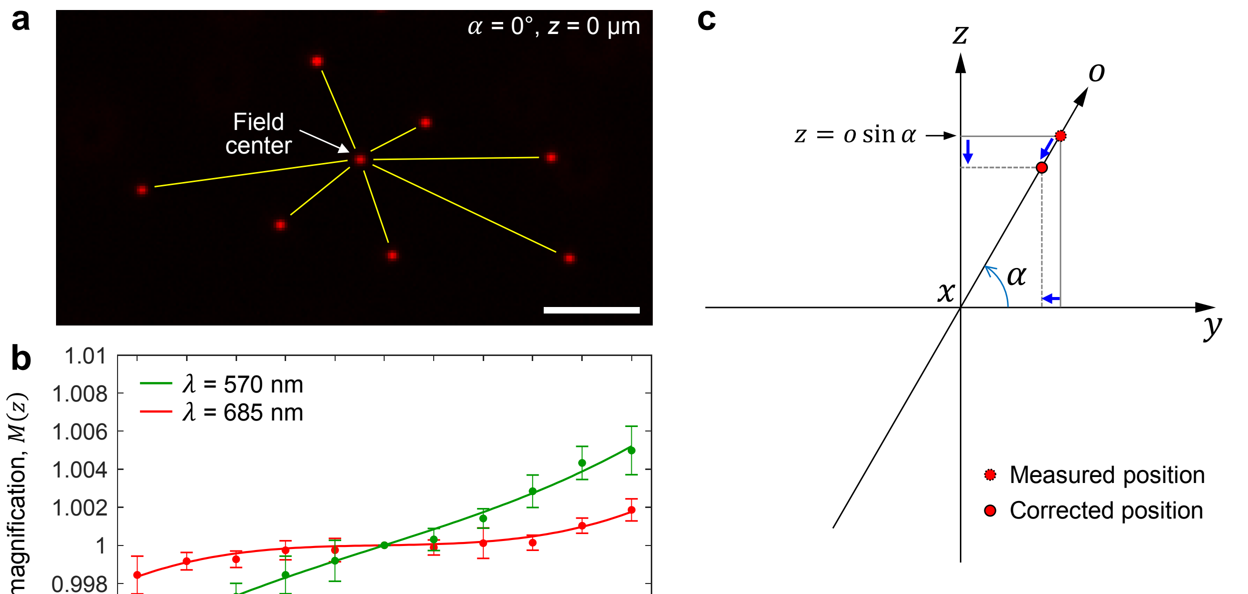


$$\begin{pmatrix} x_c \\ o_c \end{pmatrix} = \begin{bmatrix} M^{-1}(\lambda, z) & 0 \\ 0 & M^{-1}(\lambda, z) \end{bmatrix} \begin{pmatrix} x \\ o \end{pmatrix}$$

**Supplementary Figure 6 | Calibration of 3D imaging magnification.** (**a**) A 3D magnification of the setup associated with remote focusing[4] can be anisotropic (or non-uniform) due to its intrinsic residual aberrations[5] and imperfect implementation (an unavailability of a perfect set of lenses to construct the remote focusing and/or a goodness of optical alignment). To examine it, eight multi-color fluorescent beads (T7280, Invitrogen) with an emission wavelength setting of 685 nm at 0° oblique angle was imaged. The distances of 7 beads to the bead at the field center (yellow lines) were calculated over different focal positions of the beads to investigate changes in imaging magnification. Scale bar, 5 μm. (**b**) Relative change in lateral magnification through focus at two emission wavelengths. The system magnification was tuned almost uniform (near 3D isotropic) at λ = 685 nm. Then, the identical experiment in **a** was conducted for the same bead sample but with a different emission filter setting for λ = 570 nm. The setup optimized at 685 nm shows a slight ramp in magnification over depth at 570 nm. In our alignment condition, we confirmed that axial magnification through focus exhibits the same behavior with lateral magnification, and thus identified that 3D magnification is a function of wavelength and axial position ($z$). Error bars, s.d., $n$ = 7 for both wavelengths. (**c**) Schematic of magnification correction in oblique plane imaging. The two-color magnification curves in **b** were implemented here to compensate localization data. The subscript $c$ in $x$ and $o$ coordinates denotes the updated coordinates after correction.

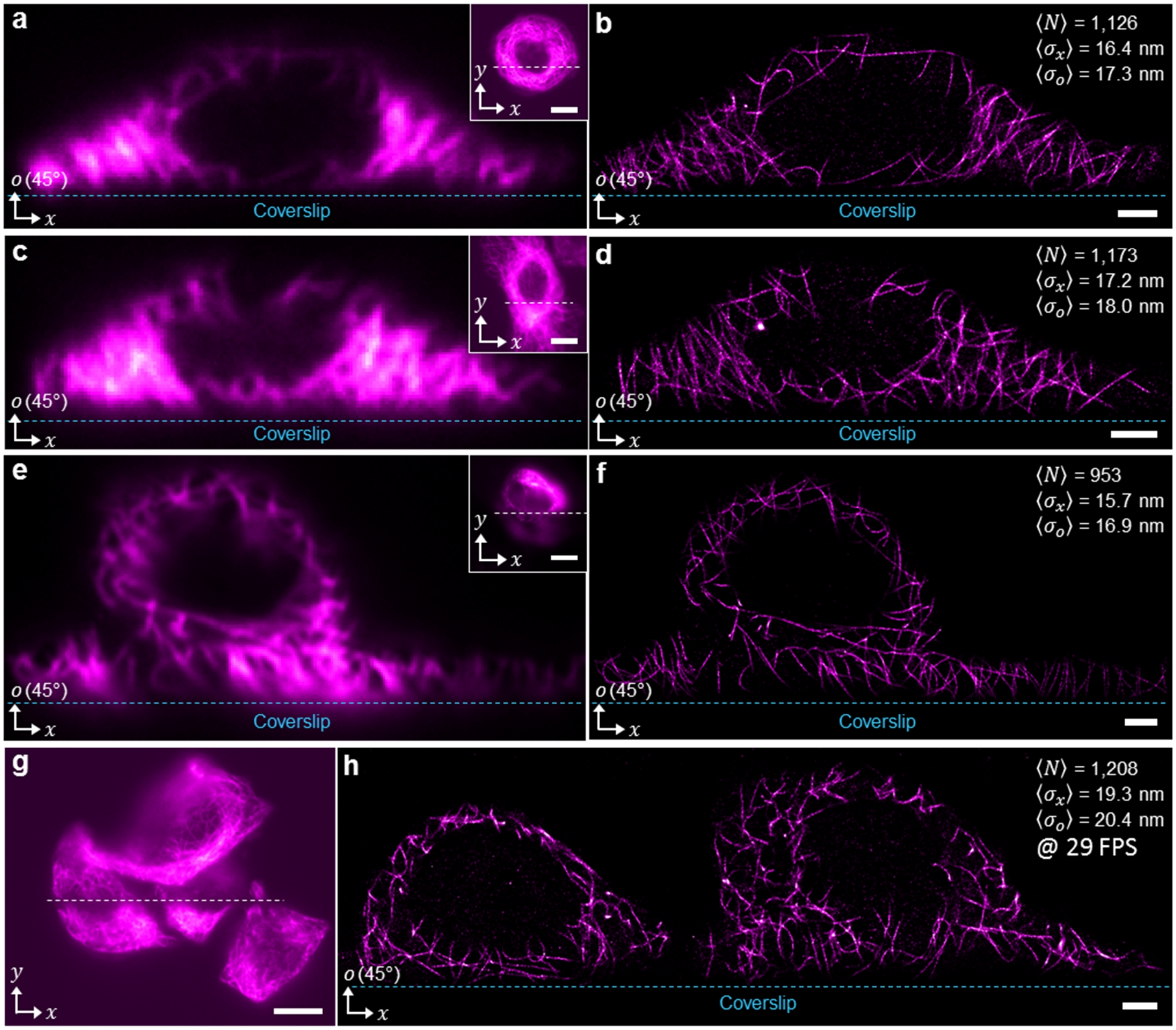


**Supplementary Figure 7 | Other 45° obSTORM images of microtubules in A549 cells.** (**a**, **c**, **e**) 45° oblique plane, diffraction-limited images of microtubules labeled with Alexa Fluor 647. The insets show the lateral plane fluorescence images with the intersection line of the two planes. (**b**, **d**, **f**) Corresponding 45° obSTORM images of **a**, **c**, and **e**. $\langle N \rangle$, the number of average photons collected; $\langle \sigma_x \rangle$ and $\langle \sigma_o \rangle$, average localization precision of dye molecules in the $x$ and $o$ directions, respectively. (**g**) Conventional fluorescence, lateral plane image of A549 cells labeled with Alexa Fluor 647. (**h**) 45° obSTORM image over three cells shown in **g**. STORM videos were taken at 50 frames per second (FPS) in **b**, **d**, and **f**. Scale bars, 2 μm (**b**, **d**, **f**, **h**) and 10 μm (Insets in **a**, **c** and **e**, and **g**).

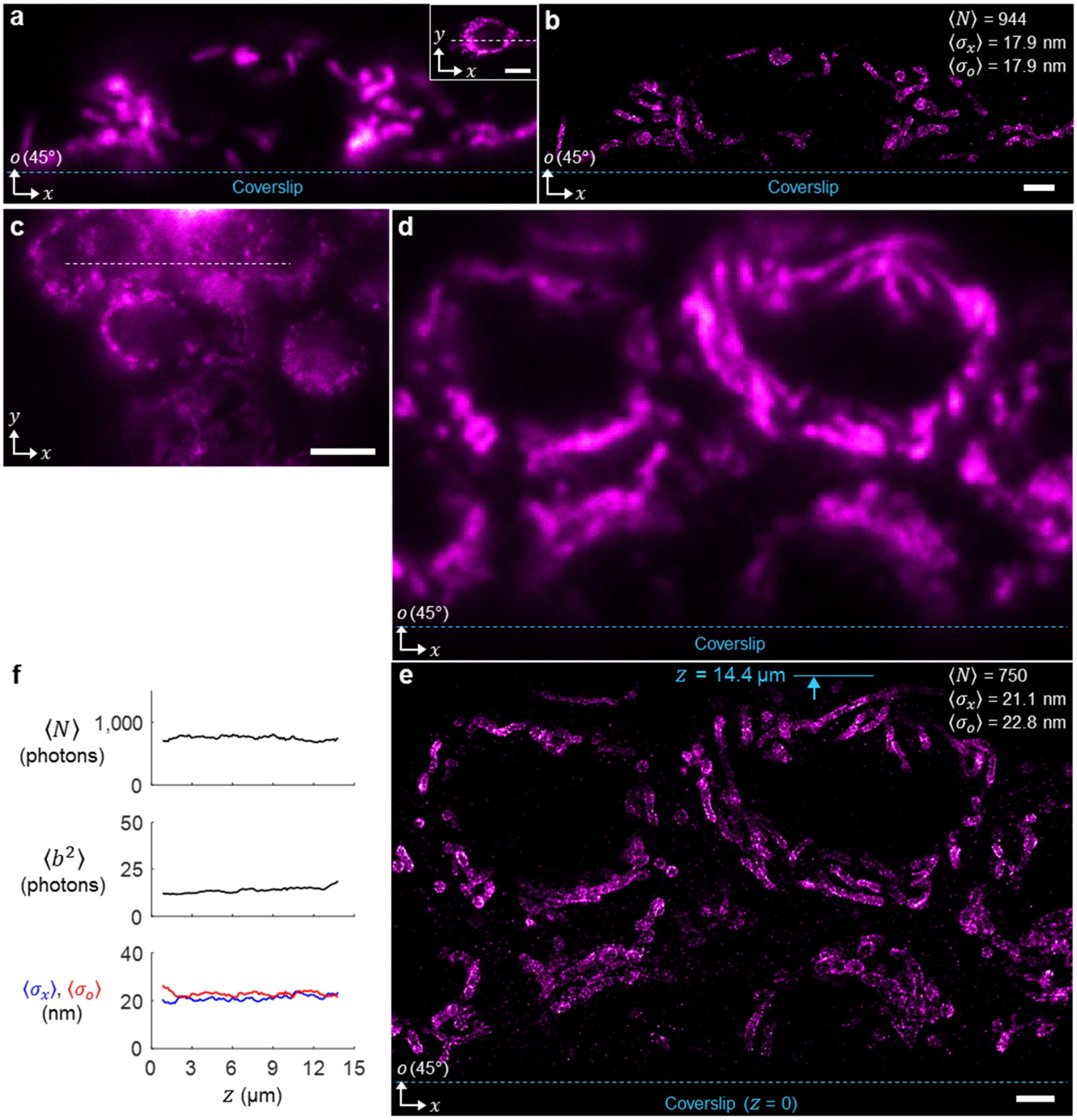


**Supplementary Figure 8 | 45° obSTORM images of mitochondria in A549 cells.** (**a**) Diffraction-limited fluorescence image of a cell along a 45° oblique plane and along the lateral plane (inset). (**b**) obSTORM super-resolution image of **a**. $\langle N \rangle$, the number of average photons collected; $\langle \sigma_x \rangle$ and $\langle \sigma_o \rangle$, average localization precision of single molecules along $x$ and $o$, respectively. (**c**) Conventional lateral plane fluorescence image of a cell cluster with a dashed line where a 45° oblique view in **d** intersects. (**d**) 45° diffraction-limited fluorescence image. (**e**) 45° obSTORM image of mitochondria in multiple cells extending up to $z$ = 14.4 μm in depth. All cells were labeled with Alexa Fluor 647. (**f**) Through-depth analysis of single molecules. $\langle b^2 \rangle$, average background photons per pixel. Scale bars, 2 μm (**b**, **e**) and 10 μm (Insets in **a** and **c**).

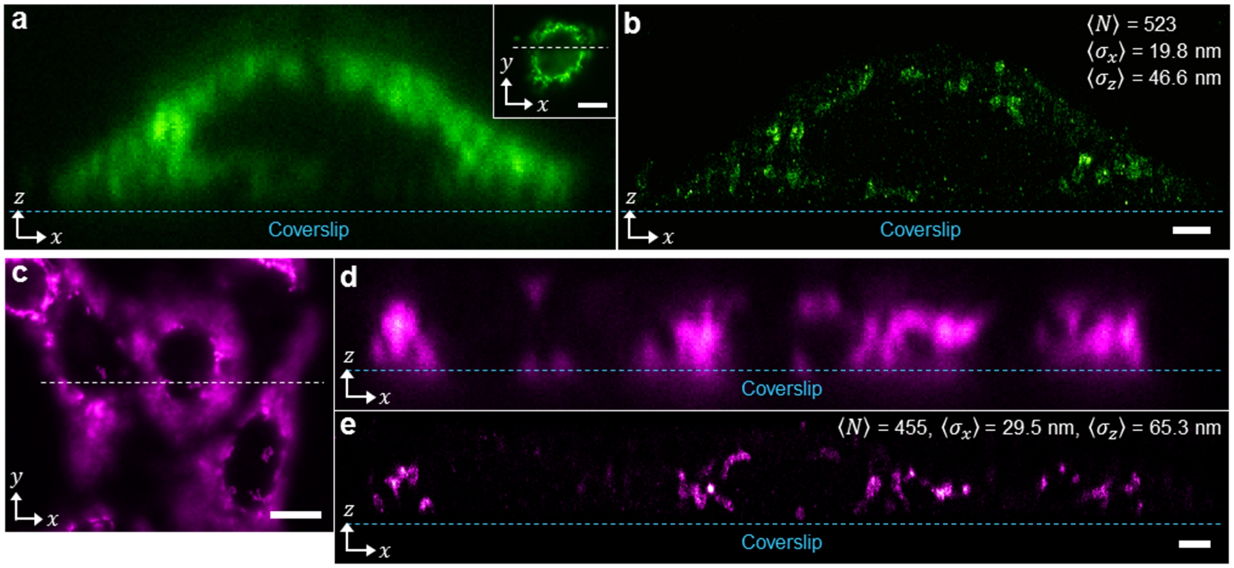


**Supplementary Figure 9 | 90° obSTORM images of mitochondria in A549 cells.** (**a**) Diffraction-limited, axial plane (α = 90°) image of mitochondria (labeled with CF568) across the intersection line in the lateral plane fluorescence image (inset). (**b**) Axial plane obSTORM image of **a**. $\langle N \rangle$, the number of average photons collected; $\langle \sigma_x \rangle$ and $\langle \sigma_z \rangle$, average localization precision of single molecules in the *x* and *z* directions, respectively. (**c**) Lateral plane conventional fluorescence image of mitochondria in multiple cells labeled with Alexa Fluor 647. (**d**) Diffraction-limited, axial plane fluorescence image across the intersection line (white dashed) in **c**. (**e**) Axial plane obSTORM image of **d**. Scale bars, 2 µm (**b**, **e**) and 10 µm (Inset in **a**, **c**).

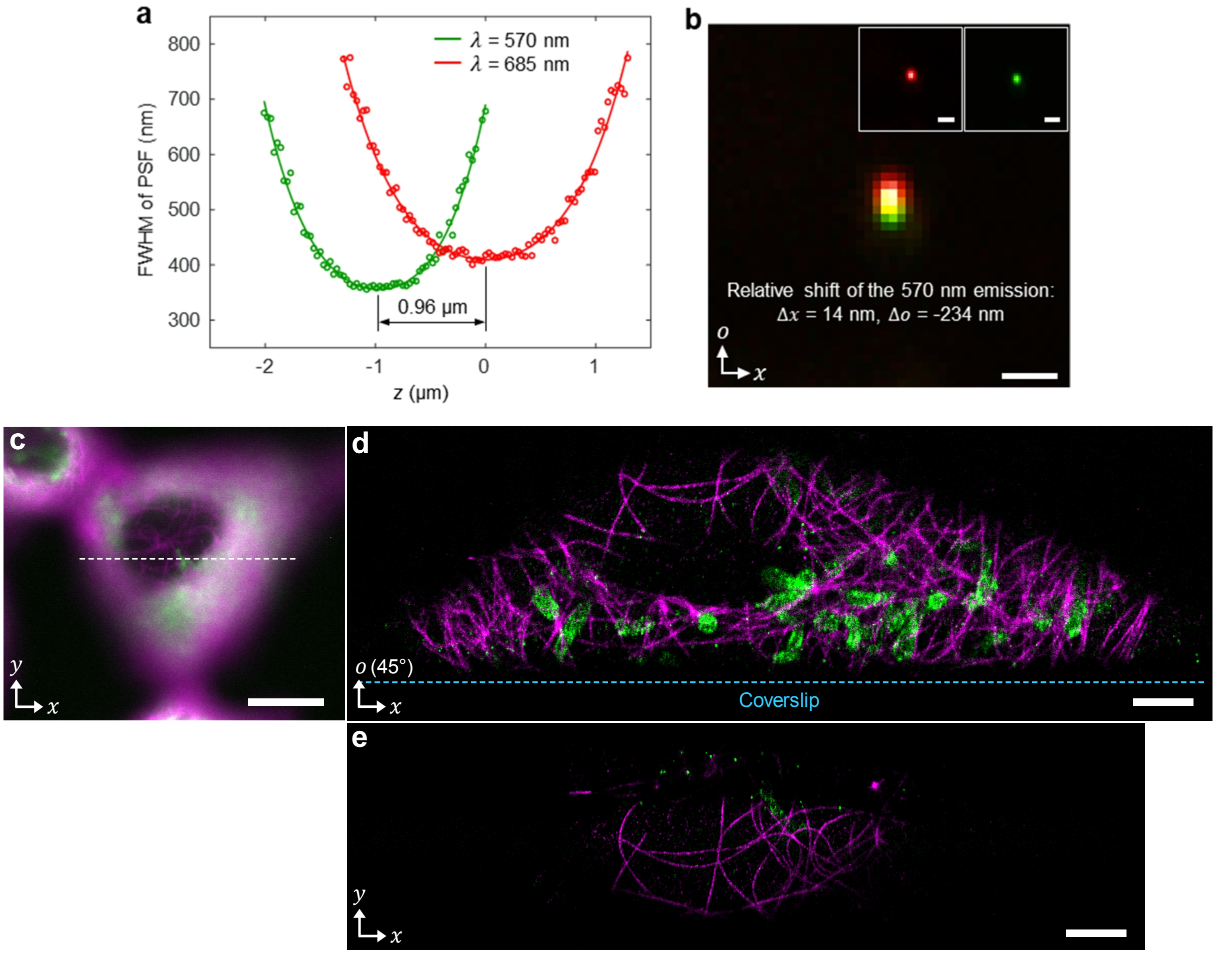


**Supplementary Figure 10 | Calibration for two-color 45° obSTORM and another example of two-color 45° obSTORM image of an A549 cell.** (**a**) Through-focus FWHM measurement of a multi-color fluorescent bead (T7280, Invitrogen) in 45° oblique plane imaging, showing the chromatic focal shift of 960 nm between 685 nm (for Alexa Fluor 647) and 570 nm (for CF568) color channels. The curves are FWHMs in the *x* direction. (**b**) A merged image of the two-color bead images at their own focus (as shown in the insets). The relative shift of the two PSFs between color channels was quantized by localization analysis. The adjustment of the focal offset affects pixel registration between the two colors primarily along the *o* axis. (**c**) Two-color conventional fluorescence image of a cell labeled with Alexa Fluor 647 for microtubules (magenta) and CF568 for mitochondria (green). (**d**) Two-color 45° obSTORM image with the calibration in **a** and **b** applied. (**e**) Two-color lateral plane STORM image of the mitochondria and microtubules along the dashed intersection in **c**. The field of view in the *y* direction is imposed mainly by the thickness and incident angle of light-sheet. Scale bars, 1 μm (all in **b**), 2 μm (**d**, **e**) and 10 μm (**c**).

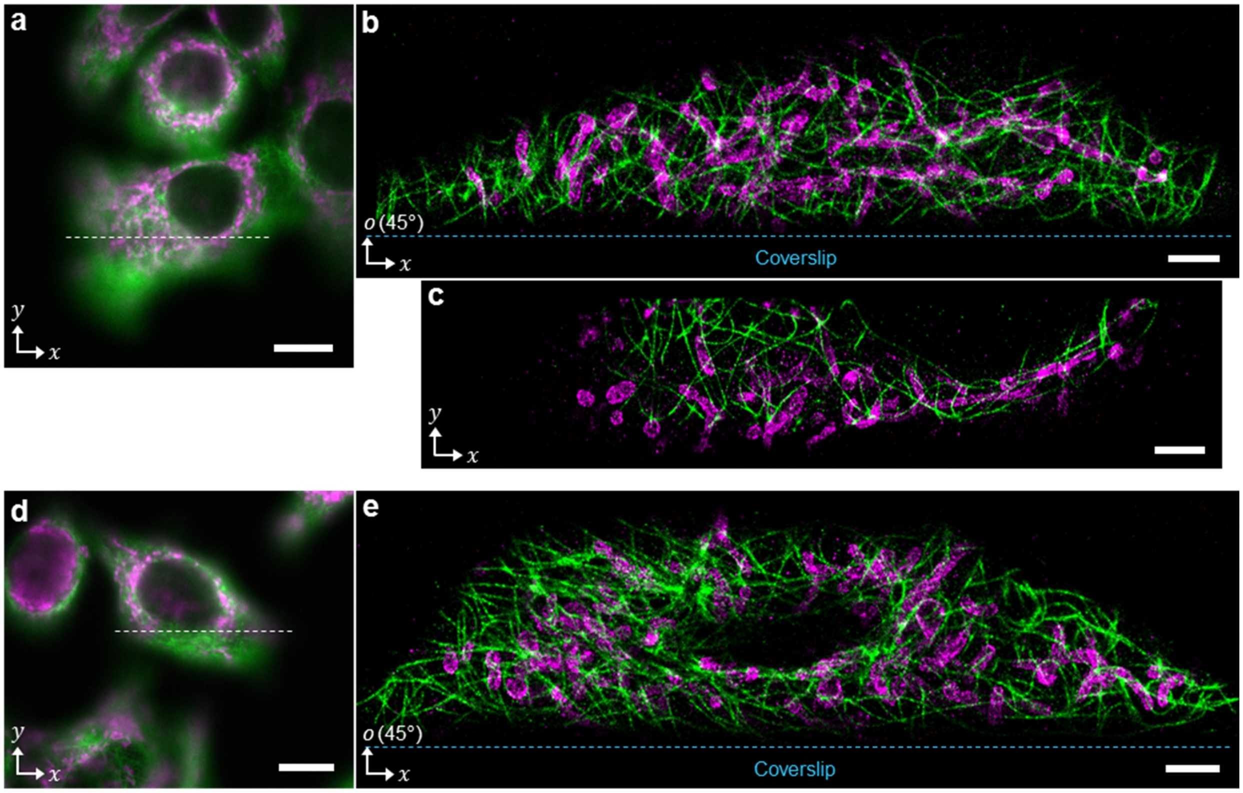


**Supplementary Figure 11 | Two other examples of two-color 45° obSTORM images with dye molecules switched in A549 cells.** (**a**) Conventional lateral plane fluorescence image. (**b**) 45° obSTORM image along the intersection in **a**. (**c**) Lateral plane STORM image near the intersection in **a** with the confined field of view (in the *y* direction) by the 45° light-sheet illumination. The field of view along *y* is imposed mainly by the thickness and incident angle of light-sheet, which appeared to be ~5.5 µm at α = 45°. Such confined field of view can vanish if a sample allows wide-field illumination. (**d**) Conventional lateral plane fluorescence image of another A549 cell. (**e**) 45° obSTORM image of the cell in **d**. Microtubules (green) and mitochondria (magenta) were labeled with CF568 and Alexa Fluor 647, respectively. Scale bars, 2 µm (**b**, **c**, **e**) and 10 µm (**a**, **d**).

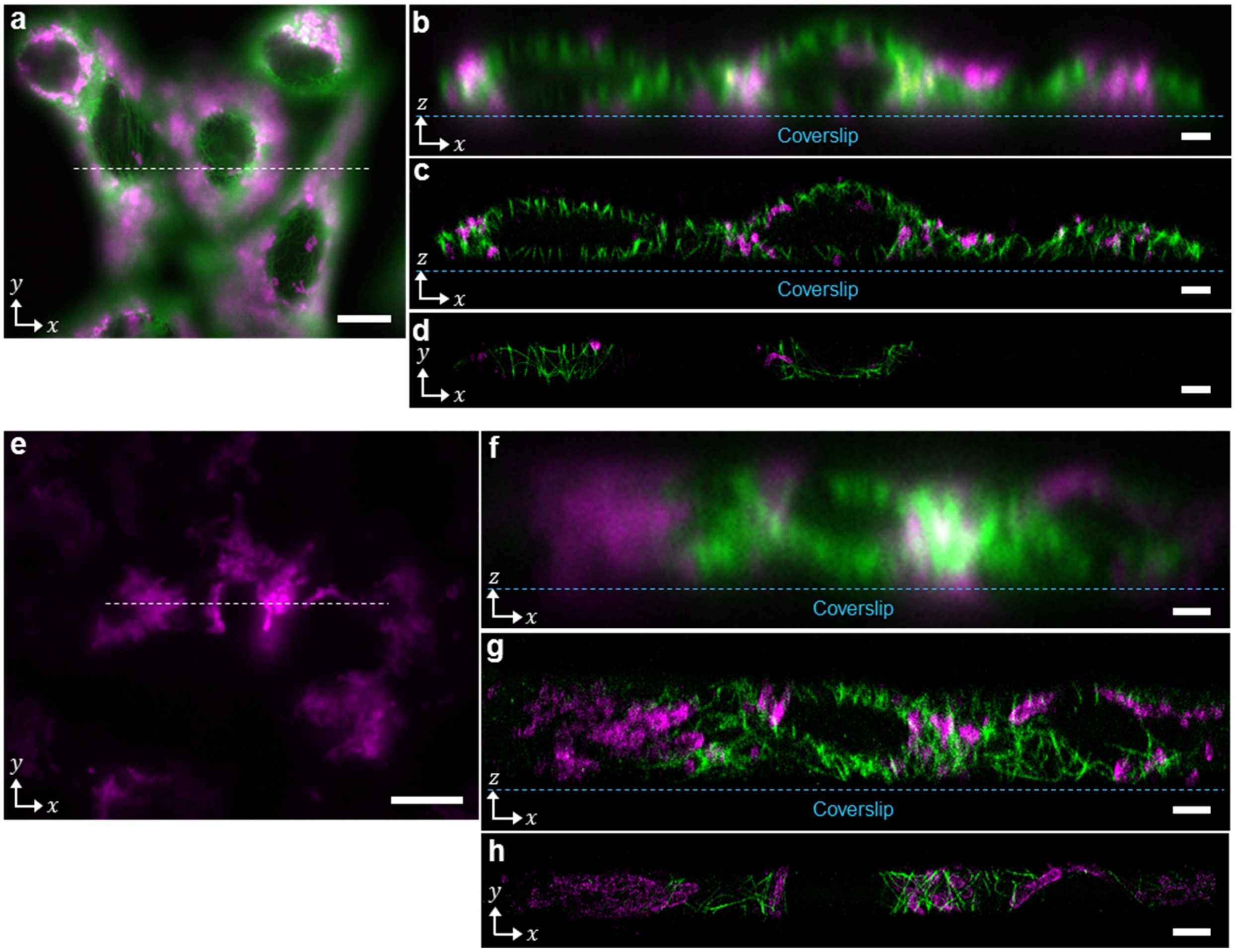


**Supplementary Figure 12 | Two-color 90° obSTORM images of microtubules and mitochondria in A549 cells.** (**a**) Conventional lateral plane fluorescence image of several cells. (**b**) Axial plane, diffraction-limited image along the dashed intersection in **a**. (**c**) 90° obSTORM image after the chromatic correction in 90° imaging mode was considered as done in **Supplementary Fig. 10a**. (**d**) Lateral plane STORM image near the intersection in **a** with the field of view in the *y* direction limited by the axial light-sheet thickness. (**e**) Conventional lateral plane fluorescence image. (**f**) 90° diffraction-limited fluorescence image along the intersection line in **e**. (**g**) Axial obSTORM image of **f**. (**h**) Lateral plane STORM image over the light-sheet thickness along the intersection line in **e**. The field of view along *y* is imposed mainly by the thickness of light-sheet and appears to be ~2.8 μm at $\alpha$ = 90°. Such confined field of view can vanish if a sample allows wide-field illumination. Microtubules and mitochondria were labeled with CF568 and Alexa Fluor 647, respectively. Scale bars, 2 μm (**b-d**, **f-h**) and 10 μm (**a**, **e**).

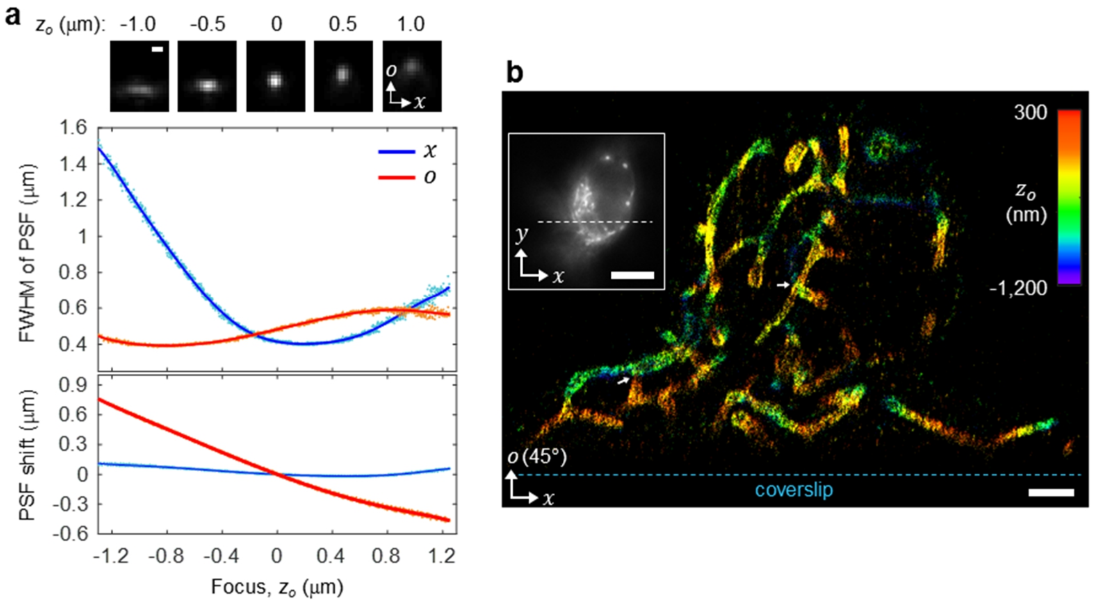


**Supplementary Figure 13 | 3D super-resolution imaging in 45° obSTORM.** (**a**) 2D PSF images at different locations of $z_o$ (normal to the oblique plane) and calibrations curves for 3D localization. To achieve 3D obSTORM on the 45° oblique plane, we placed a cylindrical lens ($f$ = 200 mm) right in front of the oblique mode EMCCD camera and measured the PSF of dark-red fluorescent beads (P7220, Invitrogen) with stepping the sample objective (O1) by 30 nm along the $z$ direction around $z$ = 0. Each bead measurement was repeated four times and the data from 10 beads was acquired. The bead images were then fitted with a 2D elliptical Gaussian function plus a constant background with the size of fitting window determined at the same criterion used in 2D localization, from which the calibration data (both FWHM and shift of PSF) were obtained and fitted with polynomial functions. As the beads were driven along the $z$ direction (rather than $z_o$) during the measurement, we multiplied $z$ by cos 45° to obtain the net amount of motion in the $z_o$ direction. Such measurement also causes an additional shift of PSF by $z$ cos 45° in the $o$ direction which is not resulted from the defocus along the $z_o$ direction, which thereby we subtracted from the raw measurement results. This made the net shift of PSF in the $o$ direction induced by the defocus along $z_o$ appear to be opposite to the measured upward shift of 2D PSF images. In 3D localization, the FWHM of PSF calibration curves was normalized as $s_i = \left(FWHM_i - FWHM_{i,\min}\right)/\left(FWHM_{i,\max} - FWHM_{i,\min}\right)$ (where $i = x, o$) to determine the $z_o$ location of each single molecule that minimizes $\left(\sum_i \left|s_{i,\text{measured}} - s_{i,\text{calib}}(z_o)\right|^2\right)^{1/2}$. Also, we quantified a 1D image distortion in the $o$ direction caused by the cylinderical lens (~11.9 %) and compensated it during image reconstruction. (**b**) 3D, 45° obSTORM image of mitochondria of an A549 cell labeled with Alexa Fluor 647. The inset shows a conventional fluorescence, lateral plane image with a dashed line where the oblique section intersects. Mitochondria at the two regions marked with white arrows clearly show that they are different mitochondrion (not connected) and cross each other. Scale bars, 500 nm (**a**), 2 μm (**b**) and 10 μm (Inset in **b**).

### Supplementary Derivation: Theoretical PSF in obSTORM

We formulate the PSF of obSTORM based on vectorial diffraction theory. The PSF can be predicted by evaluating the Debye-Wolf diffraction integral[6], $\vec{E}(\vec{x}) = -\frac{ik}{2\pi}\iint_{\Omega} \vec{E}_{\Sigma}(\vec{x}')e^{i\hat{k}_{\Sigma}\cdot\vec{x}}d\Omega$, where an electric field $\vec{E}$ (complex amplitude) in image space $\vec{x}$ is calculated if boundary electric field $\vec{E}_{\Sigma}$ and propagation vector $\hat{k}_{\Sigma}$ are given on the exit pupil $\Sigma$ described by coordinate $\vec{x}'$. $\Omega$ denotes a solid angle subtended from the exit pupil $\Sigma$ toward the geometrical focus with an infinitesimal solid angle element $d\Omega$. As demonstrated in the authors' tutorial[6] on vectorial diffraction calculation, 3×3 Jones matrix formalism assists to trace both electric field vectors and ray vectors emitted from a single molecule up to the exit pupil.

For such tracing purpose, the obSTORM layout can be simplified as below. Here, we set the Cartesian reference coordinate ($x$, $y$, $z$) such that the $z$ axis points to the right for consistent tracing[6] (different from the coordinate definition in **Fig. 1a**). We model the PBS as two linear polarizers, i.e., $\mathbb{P}(90°)$ for forward propagation and $\mathbb{P}(0°)$ for backward propagation and neglect the light reflection at the hypertense surface of the PBS. We assume that the remote mirror (M) is a perfect reflector ($\mathbb{F}_R = [1,0,0;0,-1,0;0,0,1]$) with a good accuracy[6]. While the remote mirror is supposed to rotate (or tilt) about the $y$ axis in the new coordinate, for simplicity in vectorial tracing we neglect its tilt angle in oblique plane imaging (thus, $\alpha = 0°$ for all cases). We model a fluorescent molecule as an electric dipole $\vec{p}$ located at the geometrical focus of the sample objective (OBJ1), whose far-field emission to the $\hat{k}_o$ direction can be expressed as $\vec{E}_o = (\hat{k}_o \times \vec{p}) \times \hat{k}_o$ where $\hat{k}_o = [\sin\theta_1\cos\phi_1, \ \sin\theta_1\sin\phi_1, \ \cos\theta_1]^T$ in terms of a polar angle $\theta_1$ ($\in [0,\alpha_1]$ where $\alpha_1$ is the semi-aperture angle of OBJ1) and an azimuthal angle $\phi_1$ ($\in [0,2\pi]$) in object space[6]. In remote focusing, the lateral magnification from object space to remote space is $n_1/n_r$ (a ratio of refractive indices). The magnification from remote space to image space is $M_i = f_4 f_{O2}^{-1}$. The total lateral magnification of the imaging system at $\alpha = 0°$ is $M = f_2 f_{O1}^{-1} f_4 f_3^{-1}$. We consider a semi-aperture angle ($\alpha_r$) of the remote objective (OBJ2) larger than or equal to $\alpha_1$.

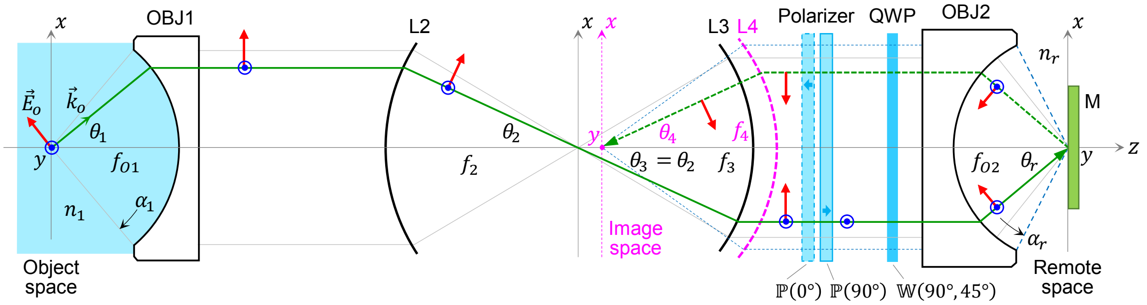


Using the Jones matrices revisited in the literature[6], the electric field vector at the exit pupil (assumed to locate right after the L4 lens) can be approximated as

$$\vec{E}_{\Sigma} = \mathbb{R}_z^{-1}(\phi_1)\mathbb{L}_4(\theta_4)\mathbb{R}_z(\phi_1)\mathbb{P}(0°)\mathbb{W}(90°,45°)\mathbb{R}_z^{-1}(\phi_1)\mathbb{L}_{OBJ_2}(\theta_r)\mathbb{R}_{y_s}^{-1}(\pi-\theta_r)\mathbb{F}_R$$
$$\times \mathbb{R}_{y_s}(\theta_r)\mathbb{R}_z(\phi_1)\mathbb{R}_z^{-1}(\pi+\phi_1)\mathbb{L}_{OBJ_2}(-\theta_r)\mathbb{R}_z(\pi+\phi_1)\mathbb{W}(90°,45°)\mathbb{P}(90°)$$

$$\times \mathbb{R}_z^{-1}(\pi+\phi_1)\mathbb{L}_3(-\theta_2)\mathbb{R}_z(\pi+\phi_1)\mathbb{R}_z^{-1}(\phi_1)\mathbb{L}_2(-\theta_2)\mathbb{L}_{OBJ_1}(-\theta_1)\mathbb{R}_z(\phi_1)\vec{E}_o$$

$$= i\sqrt{\frac{n_1\cos\theta_4}{\cos\theta_1}}\begin{bmatrix} E_{\Sigma,11} & E_{\Sigma,12} & E_{\Sigma,13} \\ E_{\Sigma,21} & E_{\Sigma,22} & E_{\Sigma,23} \\ E_{\Sigma,31} & E_{\Sigma,32} & E_{\Sigma,33} \end{bmatrix}\vec{p}$$

where

$$\begin{aligned}
E_{\Sigma,11} &= (1-\cos\theta_1)\cos\phi_1\sin\phi_1\,(\sin^2\phi_1+\cos\theta_4\cos^2\phi_1),\\
E_{\Sigma,12} &= -(\cos^2\phi_1+\cos\theta_1\sin^2\phi_1)\cdot(\sin^2\phi_1+\cos\theta_4\cos^2\phi_1),\\
E_{\Sigma,13} &= \sin\theta_1\sin\phi_1\,(\sin^2\phi_1+\cos\theta_4\cos^2\phi_1),\\
E_{\Sigma,21} &= -(1-\cos\theta_1)\cos\phi_1\sin\phi_1\,(1-\cos\theta_4)\cos\phi_1\sin\phi_1\,,\\
E_{\Sigma,22} &= (\cos^2\phi_1+\cos\theta_1\sin^2\phi_1)(1-\cos\theta_4)\cos\phi_1\sin\phi_1\,,\\
E_{\Sigma,23} &= -\sin\theta_1\sin\phi_1\,(1-\cos\theta_4)\cos\phi_1\sin\phi_1\,,\\
E_{\Sigma,31} &= -(1-\cos\theta_1)\cos\phi_1\sin\phi_1\sin\theta_4\cos\phi_1\,,\\
E_{\Sigma,32} &= (\cos^2\phi_1+\cos\theta_1\sin^2\phi_1)\sin\theta_4\cos\phi_1\,,\\
E_{\Sigma,33} &= -\sin\theta_1\sin\phi_1\sin\theta_4\cos\phi_1.
\end{aligned}$$

Using the same sequence of tracing matrices, the ray vector starting from $\hat{k}_o$ is traced at the exit pupil as $\vec{k}_\Sigma = [-\sin\theta_4\cos\phi_1\,,\ \ -\sin\theta_4\sin\phi_1\,,\ \ -\cos\theta_4]^T$. The exit pupil field for the Configuration B in **Supplementary Fig. 2** can be similarly obtained by switching $\mathbb{P}(90°)$ and $\mathbb{P}(0°)$ in the above tracing sequence.

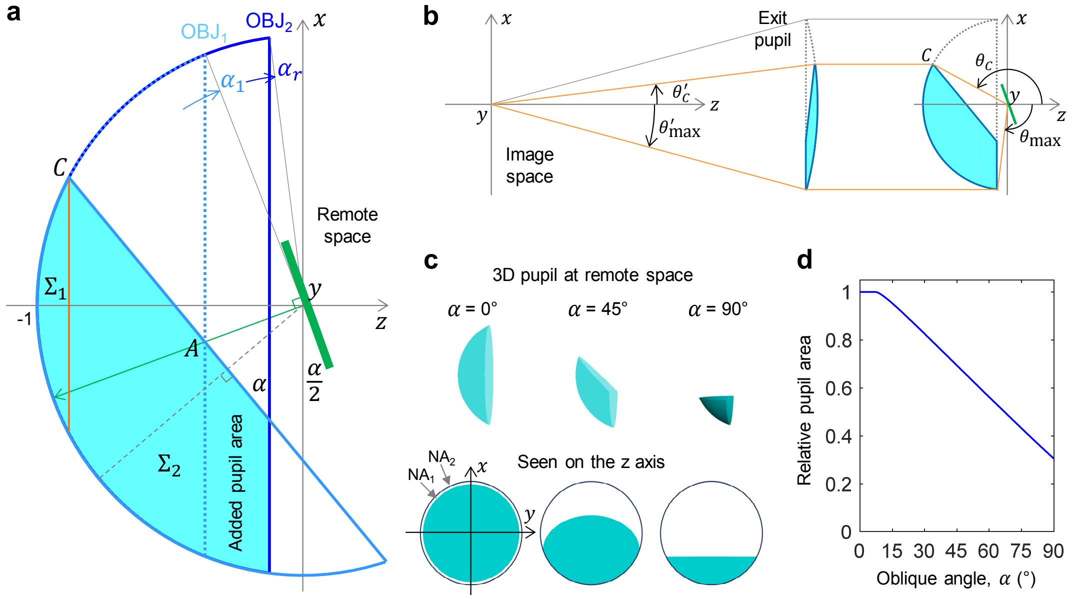


We derive a mathematical description of the 3D exit pupil in direct oblique plane imaging for a more generalized case of $\alpha_r > \alpha_1$ compared to $\alpha_1 = \alpha_r$ in earlier study[1]. As illustrated above in (**a**), the effective pupil of obSTORM can be considered in normalized remote space first. It is an overlapping area (light teal) between the reflected 3D pupil (light blue line) of the sample objective ($OBJ_1$) by the tilted mirror (green) and the 3D pupil (blue line) of the remote objective

(OBJ2). When $\alpha_r > \alpha_1$, there is an added pupil area (as labeled in (**a**) above) that increases overall signal transmission and results in brighter and smaller PSF due to the increased effective NA, thereby improving localization precision in single-molecule localization microscopy. The pupil area can be expressed in normalized spherical pupil coordinate in remote space as

$$P_{\Sigma_1,\,\mathrm{remote}}(\theta,\phi) = \begin{cases} 1, & \theta \in [\theta_C, \pi],\ \phi \in [0, 2\pi], \text{ and } x_C > 0, \\ 0, & \text{otherwise,} \end{cases}$$
$$P_{\Sigma_2,\,\mathrm{remote}}(\theta,\phi) = \begin{cases} 1, & \theta \in [\theta_{\max}, \theta_C], \phi \in [\phi_{\min}(\theta), \phi_{\max}(\theta)], \text{ and } x_C < 0, \\ 0, & \text{otherwise,} \end{cases}$$

where $\Sigma_1$ and $\Sigma_2$ denote rotationally symmetric and asymmetric parts of the pupil respectively, which are separated for mathematical convenience. $x_C$ is the $x$ coordinate of the point $C$, derived as $x_C = \sin(\alpha_1 - \alpha)$ by considering the intersection point between the circle, $x^2 + z^2 = 1$, and the line $\overline{AC}$, $z = -x \tan\alpha - \cos\alpha_1 \sec\alpha$. The symmetric part disappears when the oblique angle $\alpha$ exceeds the half-cone angle ($\alpha_1$) of OBJ1, or $x_C < 0$. The pupil boundaries can be found as

$$\theta_C = \cos^{-1} z_C = \pi - |\alpha_1 - \alpha|,$$
$$\theta_{\max} = \begin{cases} \pi - (\alpha_1 + \alpha), & \text{if } \alpha_1 + \alpha < \alpha_r, \\ \pi - \alpha_r, & \text{otherwise,} \end{cases}$$
$$\phi_{\min}(\theta) = \cos^{-1}(-\cot\alpha \cot\theta - \csc\alpha \cos\alpha_1 \csc\theta),$$
$$\phi_{\max}(\theta) = 2\pi - \phi_{\min}(\theta).$$

As the diffraction integral is evaluated on the exit pupil, the 3D pupil function derived in the remote space has to be transformed to the exit pupil function in the image space. As illustrated in (**b**) above, while the azimuthal angles in the remote and image spaces remain unchanged under such transformation, the polar angles are related with $n_r \sin(\pi - \theta_{\mathrm{remote}}) = M_i \sin\theta_{\mathrm{image}}$. Note that $\theta$ used so far is a polar angle in remote space ($\theta_{\mathrm{remote}}$) and hereafter $\theta$ represents a polar angle in image space ($\theta_{\mathrm{image}}$). Thus, the asymmetric pupil function in the exit pupil can be defined as

$$P_{\Sigma_1}(\theta,\phi) = \begin{cases} 1, & \theta \in [0, \theta'_C],\ \phi \in [0, 2\pi], \text{ and } x_C > 0, \\ 0, & \text{otherwise,} \end{cases}$$
$$P_{\Sigma_2}(\theta,\phi) = \begin{cases} 1, & \theta \in [\theta'_C, \theta'_{\max}], \phi \in [\phi_{\min}(\theta), \phi_{\max}(\theta)], \text{ and } x_C < 0, \\ 0, & \text{otherwise,} \end{cases}$$

where the exit pupil boundaries can be found as

$$\theta'_C = \sin^{-1}\left[\frac{n_r}{M_i} \cdot \sin(|\alpha_1 - \alpha|)\right],$$
$$\theta'_{\max} = \begin{cases} \sin^{-1}[n_r/M_i \cdot \sin(\alpha_1 + \alpha)], & \text{if } \alpha_1 + \alpha < \alpha_2, \\ \sin^{-1}(n_r/M_i \cdot \sin\alpha_r), & \text{otherwise,} \end{cases}$$
$$\phi_{\min}(\theta) = \cos^{-1}[\cot\alpha \cot(\sin^{-1}(M_i n_r^{-1} \sin\theta)) - \csc\alpha \cos\alpha_1 \csc(\sin^{-1}(M_i n_r^{-1} \sin\theta))],$$
$$\phi_{\max}(\theta) = 2\pi - \phi_{\min}(\theta).$$

To provide further insights into pupil shapes, the derived 3D pupils at different oblique angles for the 1.2 NA/0.95 NA system is exemplified in (**c**) above. The 0.95 NA has a slightly larger semi-aperture angle, and thus the effective pupil area at 0° (light teal) slightly underfills the exit pupil circle set by the remote objective ($\alpha_r$). At 45° and 90°, the field reflected off from the remote mirror fully fills the bottom edge of the exit pupil. We also calculated the change in a net pupil area in the remote space over oblique angle by integrating the pupil function over its boundary in (**d**) above. When $NA_1 = 1.2$ (water immersion) and $NA_2 = 0.95$, $\alpha_r$ is slightly larger than $\alpha_1$, and thus the effective pupil area does not decrease at smaller $\alpha$ up to 8°. After that, the pupil area decreases with a constant slope as the reflected light starts to be increasingly clipped at OBJ2. A collection efficiency of obSTORM over $\alpha$ would roughly follow the same trend of the reduction in the remote pupil area. If the exit pupil area is instead used for the efficiency estimation, pupil apodization induced mainly by objectives must be properly considered.

Plugging the field vector and the ray vector at the exit pupil to the Debye-Wolf integral and evaluating the surface integral on the effective pupil provide the PSF of obSTORM for a dipole source oriented along $\vec{p}$:

$$\vec{E}(\vec{x};\vec{p}) = -\frac{ik}{2\pi}\left(\int_0^{\theta_C'}\int_0^{2\pi}\vec{E}_\Sigma(\theta_1,\theta_4,\phi_1)e^{i\vec{k}_\Sigma\cdot\vec{x}}d\Omega + \int_{\theta_C'}^{\theta_{\max}'}\int_{\phi_{\min}(\theta)}^{\phi_{\max}(\theta)}\vec{E}_\Sigma(\theta_1,\theta_4,\phi_1)e^{i\vec{k}_\Sigma\cdot\vec{x}}d\Omega\right),$$

where $\vec{k}_\Sigma\cdot\vec{x} = -x\sin\theta_4\cos\phi_1 - y\sin\theta_4\sin\phi_1 - z\cos\theta_4$ and $d\Omega = \sin\theta\, d\theta d\phi$ with $\theta_4 = \theta$, $\phi_1 = \phi$ and $\theta_1 = \sin^{-1}(M_i/n_r\cdot\sin\theta_4)$. This field is calculated at the image space $\vec{x}$ ($z = 0$ for in-focus 2D PSF) and can be rescaled to the object space by a magnification factor of $M_i n_1 n_r^{-1}$ (or $M$). Note that there are two surface integrals here due to the separated pupils $\Sigma_1$ and $\Sigma_2$, and the first surface integral vanishes if $\alpha > \alpha_1$.

Finally, the intensity PSF of obSTORM, assuming the dye molecules in free-rotation, i.e., $\vec{I}(\vec{x}) = (4\pi)^{-1}\int_0^{\pi}\int_0^{2\pi}\left|\vec{E}(\vec{x};\vec{p})\right|^2\sin\theta_p\, d\theta_p d\phi_p$[6], can be derived as

$$I(\vec{x}) = \frac{1}{3}\sum_{p=1}^{3}\sum_{q=1}^{3}\left|\frac{k}{2\pi}\left(\int_0^{\theta_C'}\int_0^{2\pi}\sqrt{\frac{n_1\cos\theta_4}{\cos\theta_1}}E_{\Sigma,pq}e^{i\vec{k}_\Sigma\cdot\vec{x}}d\Omega + \int_{\theta_C'}^{\theta_{\max}'}\int_{\phi_{\min}(\theta)}^{\phi_{\max}(\theta)}\sqrt{\frac{n_1\cos\theta_4}{\cos\theta_1}}E_{\Sigma,pq}e^{i\vec{k}_\Sigma\cdot\vec{x}}d\Omega\right)\right|^2,$$

where $E_{\Sigma,pq}$ was defined during the field vector tracing.

The formulation developed here is more generalized and rigorous than our earlier derivation[1, 7] and provides sufficient physical insights into the PSF of oblique plane fluorescence microscopy. However, since the tilt angle of the remote mirror is still neglected during the field tracing, the exit pupil field approximated may be less accurate especially at higher oblique angles. More rigorous treatment of this may result in theoretical PSFs in better agreement with our experimental results.

**Supplementary Discussion: Potential improvement on resolution in obSTORM**

The resolution of obSTORM demonstrated here with 1.2 NA water immersion objective and Alexa Fluor 647 (AF647) at $\lambda$ = 685 nm is 44 nm (**Fig. 2d**) and 154 nm (**Supplementary Fig. 9e**) on 45° and 90° oblique planes respectively, which are lower than typical STORM resolution with 1.4 NA oil immersion objectives (~25 nm in $xy$, ~60 nm in $z$). The resolution in SMLM is proportional approximately to $FWHM/\sqrt{N}$ where $FWHM$ is a size of PSF of a single molecule with the $N$ number of collected photons. With this relation in mind, there are three practical ways to improve resolution as follows.

Higher NA objectives can be considered in obSTORM. For example, with the 1.27 NA (water)/1.49 NA (oil) configuration, an expected improvement is 1.7-fold (**Supplementary Fig. 3**). While aqueous buffer in SMLM may be beneficial for live samples, sample indices elevated to near-oil (~1.50)[8] have already been reported and can still be useful. In this regard, higher NA (1.45-1.49) oil immersion objectives can be used to construct the remote focusing layout, making obSTORM's PSF smaller and improving collection efficiency due to increased pupil areas with higher cone angles (**Supplementary Fig. 3** and **Supplementary Derivation**). Thus, the improvement in resolution at $\alpha$ = 90° (the worst resolution case) compared to the current 1.2 NA/0.95 NA system is expected to be 2-fold (1.45 NA/1.49 NA) and 2.6-fold (1.49 NA/1.49 NA) as estimated in **Supplementary Fig. 3**.

An up-to-date scientific complementary metal-oxide-semiconductor (sCMOS) camera can improve the localization precision by 1.4-fold[9]. Recent back-illuminated sCMOS cameras (Prime 95B, Photometrics) have a comparable quantum efficiency with back-illuminated EMCCD cameras commonly used in SMLM. They are free from the excess noise factor of ~2 induced in EMCCD cameras during electron multiplication processes[10]. The 1.4-fold improvement of localization precision (and thus resolution) was experimentally proved[9].

Chemistry plays an important role on switching organic dye molecules in STORM. Adding a triplet-state quencher such as cyclooctatetraene in typical STORM buffers is known to increase the photo-stability of AF647 and thus yield ~3.5-fold increased photons without any side effects[11]. This simply modified buffer itself independently can result in 1.87-fold improvement in resolution over all oblique angles.

By using the advanced sCMOS camera and the improved STORM buffer together, an expected resolution in the current 1.2 NA/0.95 NA system is 2.6-fold. This means that the resolution at 45° and 90° obSTORM with AF647 will be 17 nm and 59 nm respectively, which are comparable to conventional STORM resolution achieved with 1.4 NA (oil) objectives. Thus, <60 nm resolution may potentially be possible in the current water immersion system throughout all oblique angles. A further improvement enabled by high-end commercial objectives may allow higher resolution in 90° obSTORM (the worst case) with AF647 to be ~31 nm with 1.27 NA (water)/1.49 NA (oil) objectives and ~23 nm with 1.49 NA/1.49 NA objectives.

### References for the Supplementary Information


1. Kim, J., Li, T., Wang, Y. & Zhang, X. Vectorial point spread function and optical transfer function in oblique plane imaging. *Opt Express* **22**, 11140-11151 (2014).

2. Anselmi, F., Ventalon, C., Begue, A., Ogden, D. & Emiliani, V. Three-dimensional imaging and photostimulation by remote-focusing and holographic light patterning. *Proc. Natl. Acad. Sci. USA* **108**, 19504-19509 (2011).

3. Cella Zanacchi, F. et al. Live-cell 3D super-resolution imaging in thick biological samples. *Nat Meth* **8**, 1047-1049 (2011).

4. Botcherby, E.J., Juskaitis, R., Booth, M.J. & Wilson, T. Aberration-free optical refocusing in high numerical aperture microscopy. *Opt Lett* **32**, 2007-2009 (2007).

5. Botcherby, E.J., Juskaitis, R., Booth, M.J. & Wilson, T. An optical technique for remote focusing in microscopy. *Opt Commun* **281**, 880-887 (2008).

6. Kim, J., Wang, Y. & Zhang, X. Calculation of vectorial diffraction in optical systems. *J. Opt. Soc. Am. A* **35**, 526-535 (2018).

7. Li, T. et al. Axial Plane Optical Microscopy. *Scientific Reports* **4**, 1-6 (2014).

8. Olivier, N., Keller, D., Rajan, V.S., Gönczy, P. & Manley, S. Simple buffers for 3D STORM microscopy. *Biomedical Optics Express* **4**, 885-899 (2013).

9. Camera Comparison: Prime 95B™ Scientific CMOS and EMCCD (https://www.photometrics.com/resources/technotes/pdfs/CameraComparison-Prime95B-sCMOS-and-EMCCD-TechNote.pdf; 2018).

10. Hynecek, J. & Nishiwaki, T. Excess noise and other important characteristics of low light level imaging using charge multiplying CCDs. *IEEE Transactions on Electron Devices* **50**, 239-245 (2003).

11. Olivier, N., Keller, D., Gönczy, P. & Manley, S. Resolution Doubling in 3D-STORM Imaging through Improved Buffers. *PLOS ONE* **8**, e69004 (2013).
